# Focal neurostimulation of propagating calcium waves in human neural cells using megahertz-range single-pulse focused ultrasound

**DOI:** 10.1101/2025.07.02.662731

**Authors:** Tom Aubier, Ivan M. Suarez-Castellanos, Magali Perier, Gilles Huberfeld, Alexandre Carpentier, W. Apoutou N’Djin

**Affiliations:** LabTAU, INSERM, Centre Léon Bérard, Université Claude Bernard Lyon 1, F-69003, Lyon, France; Department of Neurology, Hôpital Fondation Adolphe de Rothschild, 75019 Paris, France; Université Paris Cité, Institute of Psychiatry and Neuroscience of Paris (IPNP), INSERM U1266, Neuronal Signaling in Epilepsy and Glioma, 75014 Paris, France; Sorbonne Université, AP-HP, Hôpitaux Universitaires La Pitié Salpêtrière - Charles Foix, Service de Neurochirurgie, Paris, France

**Author notes:** **Corresponding authors:**, &.

**Keywords:** Single-pulse Focused Ultrasound, FUS, Focal Neurostimulation, Neuromodulation, Human Neural Cells, Calcium waves, In vitro, Focal responses, Propagative responses, Wearable Transdural Ultrasound Intracranial Implant

## Abstract

Focused ultrasound (FUS) offers a promising neurostimulation technique that addresses the trade-off between invasiveness and spatial selectivity found in conventional electromagnetic-based methods. Evidence supporting its theoretical selectivity and comprehensive descriptions of the underlying biophysical mechanisms remain insufficient to ensure adequate control and safety. Based on observations of causal, focal, and propagating calcium waves in human neural cells in vitro evoked using single-pulse megahertz-range FUS, a transdural FUS approach relying on an intracranial implant is proposed to overcome frequency limitations associated with transcranial ultrasound. Using an extended parameter space (0.7 −8 MHz), calcium waves were shown to be selectively elicited by cavitation or acoustic radiation force. Cavitation, predominant at frequencies under 5 MHz, induced dispersed and unpredictable responses that could lead to cell damage. FUS radiation force however, predominant at frequencies over 5 MHz, elicited focal and predictable responses, without affecting cell viability. FUS-evoked intercellular calcium waves were shown to propagate across the surrounding neural network via intracellular and extracellular pathways driven by calcium amplification mechanisms. Collectively, these findings lay the foundation for developing wearable neurostimulation approaches to manage chronic neurological conditions.

## 1 Introduction

Neurological disorders represent the leading cause of disability and stands as the secondary cause of deaths worldwide [2]. Acting directly on the behavior of affected neural structures, by eliciting or modulating their activity can improve the symptoms of patients suffering from these conditions. With an estimated 14 000 *Deep Brain Stimulation* (DBS) devices implanted in 2022 [3, 4], this modality has become a gold standard by locally imposing electrical currents to affect the activity of targeted brain structures. However, the risks, costs and technical challenges associated with electrodes implanted in brain tissues [5] have indeed prevented its widespread adoption. Alternative and non-invasive modalities such as *Transcranial Magnetic Stimulation* (TMS) or *transcranial Direct Current Stimulation* (tDCS) enable brain regions to be reached transcranially. The diffuse nature of electrical and magnetic fields however, imposes strong limitations on the energy levels deposited on specific, and especially deep neural structures, thereby weakening the magnitude of the elicited immediate biological effects. Following the definitions proposed by Gavrilov’s foundational work [6], specific terminologies can be used to clarify the outcomes of the various modalities. While *neuromodulation* refers to the process of influencing the inherent activity of a neural structure by modifying its state oftentimes using low energy but prolonged exposures to the stimuli - *neurostimulation* describes the initiation of neural activity consequent to a short and rarely repeated stimulus. Finally, it should be noted that all aforementioned modalities are not suited to patients in equivalent pathological situations. Although invasive, DBS relies on wearable implants capable of managing chronic diseases requiring day-to-day management at home. On the other hand, despite being fully non-invasive and thus less aggressive procedures for patients, techniques such as TMS or tDCS cannot be used without involving neurological experts and neuronavigation systems. To envision the application of these methods to a wide range of therapeutic needs, it is therefore apparent that fundamental research is needed to improve neurostimulation / neuromodulation modalities. This involves on one hand technical progress towards minimally-invasive techniques capable of targeting precise, potentially deep brain regions. And on the other hand, requires refining the control over biological outcomes, achieved through a better understanding of the impact of energy deposition on neural structures.

First documented in 1929 [7], it is now recognized that ultrasound can affect the activity of neural structures. The propagation properties inherent to this mechanical wave, already widely exploited in medical imaging and therapeutic applications, make *Focused UltraSound* (FUS) an appealing minimally-invasive and yet targeted potential neurostimulation / modulation modalities.

From a technical standpoint, a number of strategies have been explored to deliver ultrasound in the brain (for thermal ablation, drug delivery and neuromodulation) covering a large range of *spatial selectivity, targeting depth* and *invasiveness*: *1*. single-element transducer-based transcranial FUS (TUS) techniques to target cortical structures [10], *2*. multi-element [11], or lens-equipped TUS systems to reach deeper brain regions [12–14], *3*. transdural ultrasound implants to eliminate low-frequency limitations imposed by the skull[15, 16]. Various temporal sequences have also been explored for ultrasound neurostimulation / modulation. All proposed strategies involve low-energy FUS sequences to safely affect functional activities while avoiding irreversible damages to the brain. While most of them include long-lasting, low-intensity repetitive-pulse FUS sequences, few studies have also explored the effects of single pulses of higher intensities. Ranging from brief, high-intensity single pulses [17, 18] to long-lasting, low-intensity repetitive-pulse sequences [19, 20], there exists a broad parameter space yielding a potentially equally diverse set of elicited biological responses. In the spirit of the complexity gradation of TMS temporal sequences [21] and given the complex nature of the problem, there is a clear interest to better understand the effects of simple single-pulse FUS exposures on neural structures, before considering their safe application in single-or more complex repetitive-pulse protocols.

On the basis of ultrasound bioeffect studies that started in the first half of the 20^th^ century [22–24], a collective research effort currently consists in exploring interactions between ultrasound-induced physical perturbations and the function of the micro-structures supporting neural activity. Among others, temperature increase [25], intra-membrane cavitation [26], or displacement-mediated effects such as capacitive effects [27, 28] and mechanosensitive ion channels activation [29–31], have been proposed. While these propositions give insights as to how ultrasound might immediately trigger or modulate the function of neural structure sub-units, there is no consensus on their implication in the mid-to long-term effects observed at higher functional levels in *in vivo* studies [10, 32]. Studies have been initiated to identify the cellular processes activated or inhibited after delivery of ultrasound stimulus. *In vivo* microdialysis studies [19, 33] identified variations in extracellular GABA or Dopamine and Serotonin, arising tens of minutes after sonication periods. Similar observations were recently made using magnetic resonance spectroscopy in humans [34]. *In vitro* works, conducted using micro-electrode arrays or calcium fluorescence imaging [18, 35, 36] are also starting to uncover the involvement of specific neurotransmitter-based signaling pathways and interactions between different neural cell types. A recent study notably reported modulation of glutamate-mediated current flow lasting for up to 12 hours [37].

Overall, current challenges consist in finding stimulation parameters and associated predominant mechanisms allowing for safe, repeatable and targeted modulation of neuronal functions. Ultrasound frequencies in the megahertz range, which have been under-investigated due to limitations associated with the passage of the skull in the context of fully non-invasive approaches, should also be considered to shed light on the therapeutic potential of minimally-invasive ultrasound delivery approaches bypassing these limitations. Additionally, more complete descriptions of temporally and spatially remote effects, occurring after stimulations and at a distance from targeted regions, would also be of great help in mitigating potential adverse effects.

Here, we report on a first attempt at identifying and characterizing intercellular signaling pathways causally mobilized by spatiotemporally-selective FUS neurostimulation. This research was conducted in the prospect of developing transdural FUS brain stimulation implants in the context of the management of chronic neurological diseases. In deed, like DBS, the wearable nature of this approach makes it particularly suited for daily treatments while remaining non-invasive for brain tissues and overcoming frequency limitations encountered with TUS approaches (Fig. 1-a). Given the aforementioned challenges, we focused our attention on the effect of FUS on the neural structure only, and investigated the relevance and viability of using higher FUS frequencies to further improve the spatial selectivity of neurostimulations. *Single-pulse* stimulation scheme was chosen in an effort to gain insights on the causal biological response following the fundamental building block of any complex *multiple-pulse* ultrasound sequence. We conducted experiments on 2D *in vitro* human neural cell cultures composed of interconnected neuronal and glial cells (Fig. 1-b-c). The calcium ion (Ca^2+^) being involved in a wide range of neural signaling processes, we relied on Ca^2+^ fluorescence microscopy to assess cellular activity at the network scale. The spatial-temporal dynamics of single-pulse FUS-evoked Ca^2+^ neural responses were investigated to provide insights on the biophysical and neurobiological mechanisms underlying FUS neurostimulation. Given the frequency-dependence of the physical processes involved in ultrasound-to-biological-tissues interactions, our investigations were conducted over a wide range of FUS frequencies and allowed - through the study of the spatiotemporal dynamics of Ca^2+^ responses - the identification of two stimulation regimes induced either by the FUS field directly emitted by the transducer in a controllable manner or indirectly through stochastic FUS-induced cavitation. This strategy demonstrated that the predominance of biophysical ultrasound mechanisms inducing Ca^2+^ activity can be modulated in FUS neurostimulation procedures through the choice of central frequencies using a unique FUS implantable prototype. Finally, through systematic and quantitative assessment of the spatiotemporal dynamics of evoked responses, pharmacological blockade procedures and alterations to extracellular medium states, we identified the role of specific cell signaling pathways involved in megahertz-range single-pulse FUS stimulations.

**Figure 1.**
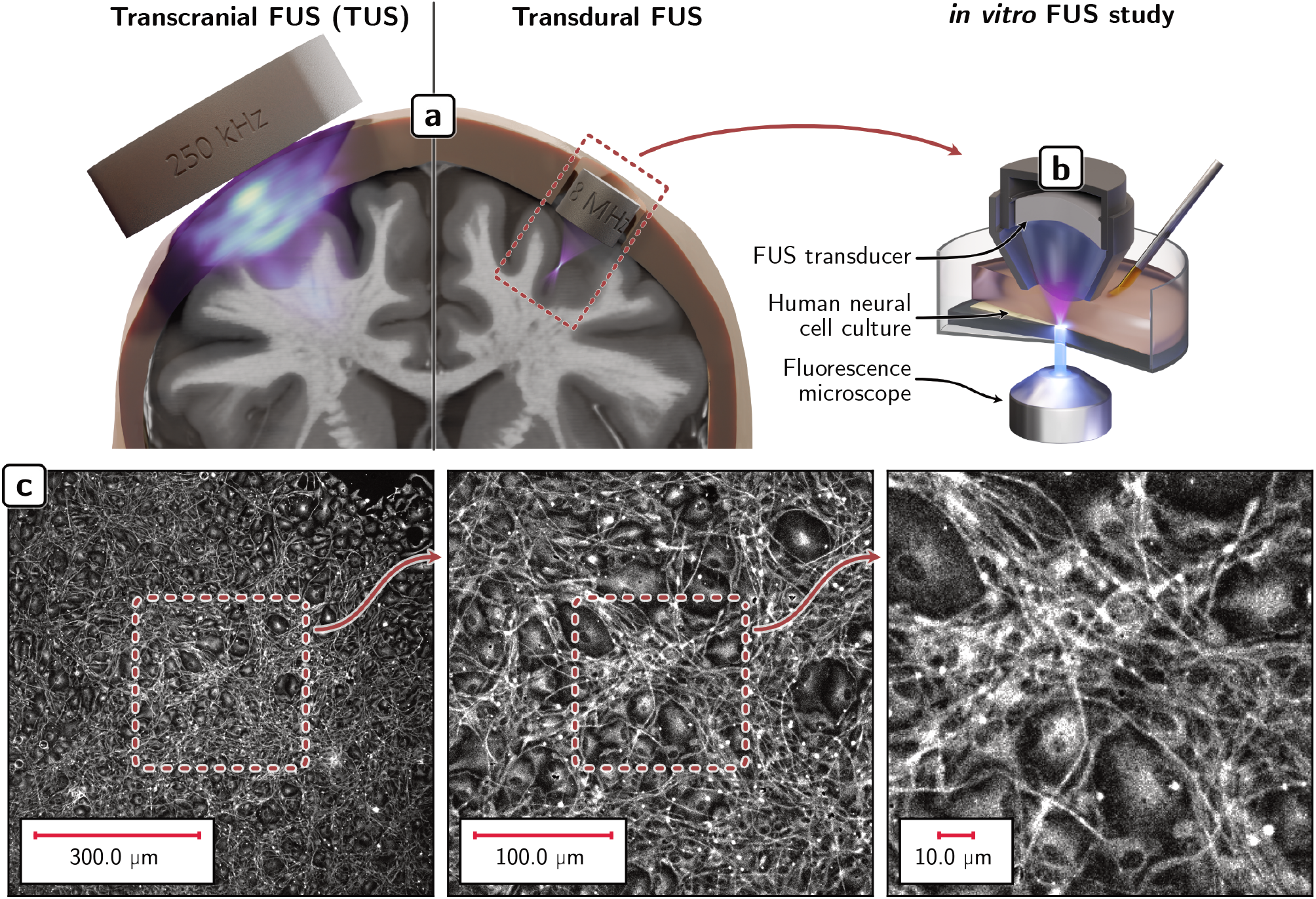
Single-pulse focused ultrasound (FUS) neurostimulation investigations on an in vitro human neural cell model in the context of an transdural intracranial FUS approach. (a) Illustration^1^ of a FUS stimulation approach based on an transdural FUS cranial implant, enabling highly selective and chronic stimulation schemes. (b) Schematic representation of the in vitro FUS neurostimulation set-up, enabling simultaneous assessment of cellular activity across a large part of the cellular network. The set-up consists in a Ca^2+^ fluorescence microscopy imaging platform equipped with a custom-made single-element FUS transducer. (c) 2D in vitro human neural cell cultures composed of interconnected neuronal and glial cells. Immortalized human neural progenitors (ReNcell VM, Merck Millipore) were cultured and differentiated according to the protocol described in [1].

## 2 Results

### 2.1 Single-pulse FUS neurostimulation of focal and propagating Ca^2+^ responses

Ultrasound experiments were conducted *in vitro* using a unique mono-element FUS transducer (15 mm in diameter and focal distance) compatible with the proposed transdural FUS intracranial implantable approach. Escalation of dose performed at frequencies ranging from 0.73 to 8.14 MHz allowed the identification of an elementary brick of neurostimulation in the form of a 400 µs single-pulse FUS sonication (Fig. 2-a-c). In these conditions, low FUS pressures (*p*_sar_ ≈ 0.1 MPa) were not able to elicit measurable Ca^2+^ activities. Increased FUS pressures (0.1 MPa *< p*_sar_ *<* 1 MPa) could elicit short-term (*<* 1 s-long) Ca^2+^ responses restricted to individual, randomly distributed cell bodies, with no evident spatiotemporal coherence. This response regime was classified as *sparse* (Fig. 2-g). No clear intercellular Ca^2+^ signaling was identified at this stage. Initiation of such *propagating* Ca^2+^ responses was shown possible at higher acoustic pressures (*p*_sar_ ≈ 8 MPa). However, using sub-megahertz FUS frequencies at these pressure levels drastically increased the occurrence of large-scale cavitation events, associated to *dispersed* responses and visible cell damage. Leveraging higher FUS frequencies appeared as a viable way to reduce or eliminate cell damage, while initiating *focal* responses with refined spatial targeting and improved repeatability (Fig. 2-d-f).

**Figure 2.**
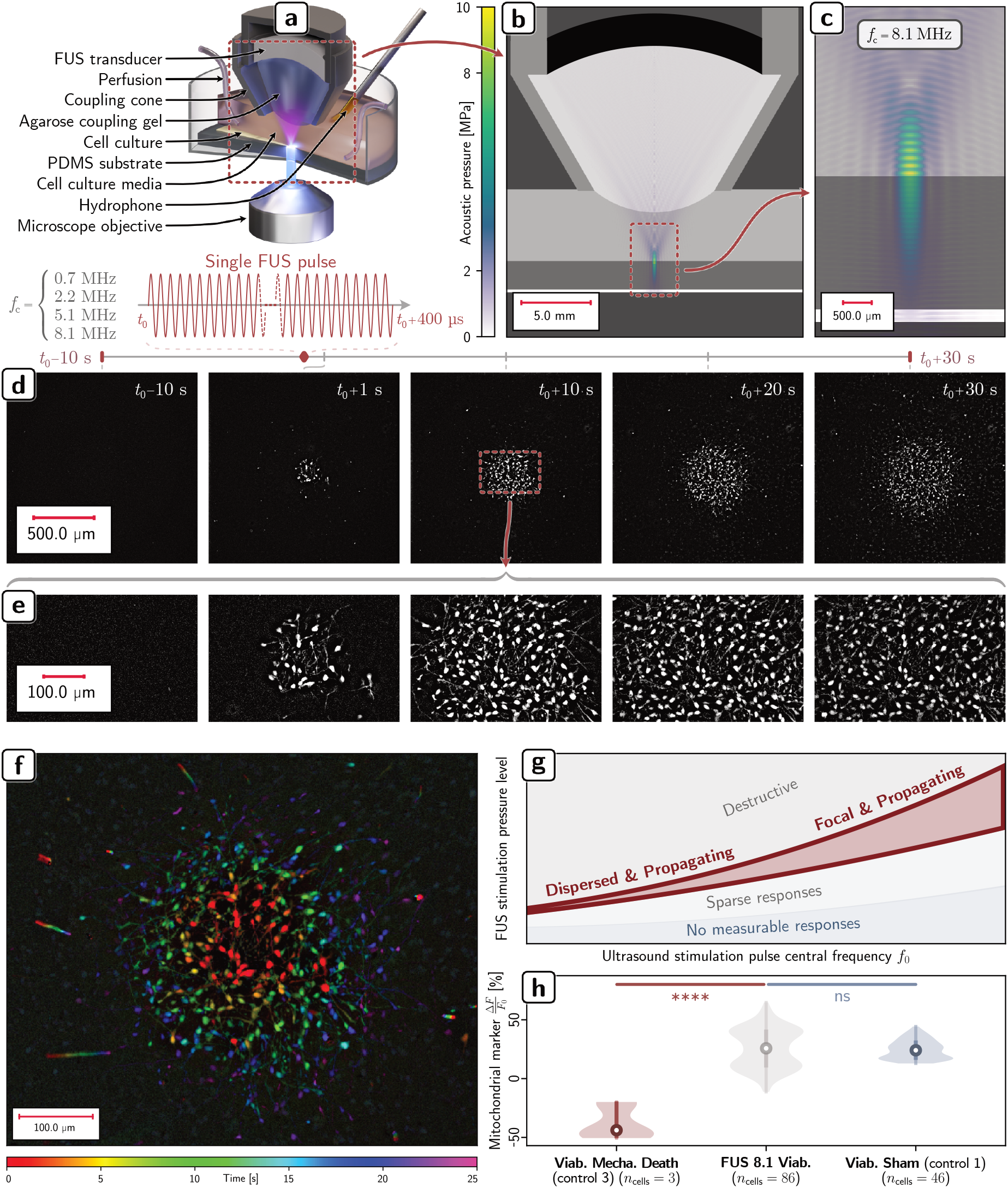
Immediate focal and delayed propagating Ca^2+^ waves induced by single-pulse FUS in human neural cells in vitro. Feasibility and safety. **(a)** Schematic depiction of the in vitro FUS neurostimulation set-up, enabling simultaneous assessment of cellular Ca^2+^ activity across a large portion of the cellular network (fluorescence microscopy imaging), as well as passive detection of FUS-induced cavitation activity (hydrophone measurements). Single-pulse FUS protocols of 400 µs were investigated accros FUS frequencies ranging from 0.73 to 8.14 MHz delivered using a unique implantable transducer prototype. **(b)** Numerical simulation of the acoustic pressure field imposed by the FUS transducer prototype at 8.14 MHz in the context of in vitro neural cell culture experiments presented in this work. **(c)** Close up of the numerically modeled focal spot for a 400 µs FUS sonication at 8.14 MHz. **(d)** Feasibility of evoking immediate focal and delayed propagating Ca^2+^ responses at the network scale with 400 µs 8.14 MHz single-pulse FUS. Fluorescence microscopy images cover 10 s pre-FUS to 30 s post-FUS exposures (**FUS 8.1** group). **(e)** Close up view of responding cells. **(f)** Compound representation of a focal and propagating Ca^2+^ intercellular response elicited by FUS. Compound images were formed by summing the stack of fluorescence time differences to which an RGB value was assigned based on the color map shown in the lower part of the figure. This representation of time is used throughout the article. **(g)** Schematic representation of response types identified experimentally through exploration the frequency vs. pressure level FUS parameter space. Results presented in this work are limited to responses falling in the focal and propagating responses category. **(h)** Safety of single-pulse FUS neurostimulation evaluated through mitochondrial viability tests (see **FUS 8.1 Viab**. vs. **Viab. Sham** (control 1) and **Viab. Mecha. Death** (control 2) groups).

The feasibility of evoking causal, focal and propagating Ca^2+^ responses repeatedly using 400 µs single-pulse FUS stimulation protocols at high-frequencies (*>*1 MHz) was demonstrated. Less than 1 s after the initiation of 8.14 MHz, 400 µs-long single-pulse FUS stimulations (*p*_sar_ = 8 ±1 MPa, *I*_sapa_ = 5 ± 1 kW.cm^−2^), focal, strong and sustained Ca^2+^ concentration elevations were observed for cells located within the targeted region. As shown on Figure 2-d-f, elicited Ca^2+^ activity led to a slow wave propagating omnidirectionally throughout the network, at an average velocity of *v*_[0,5]*s*_ = 24 ± 4 µm.s^−1^. The response expansion then continued to slow down *v*_[20,25]*s*_ = 5±1 µm.s^−1^ before reaching a peak extent in the order of 800 µm, 30 s after the stimulation. This *focal and propagating* response regime was more reproducible and controllable (78% success rate over 18 trials) than the *dispersed* regime, which prompted the detailed analysis of the biophysical mechanisms associated to high-frequency (8.14 MHz) high-pressure (*p*_sar_ ≈ 8 MHz) and short (400 µs) single-pulse FUS stimulation protocols (*Suppl. Video FUS 8*.*1*).

Assessment of mitochondrial viability over a 10 min-long period following 8.14 MHz single-pulse FUS showed no statistically significant differences (*p* = 0.235) between FUS and sham groups (see **FUS 8.1 Viab**. vs **Viab. Sham** (control 1) groups). In contrast, control tests consisting in cell membrane perforation performed with a patch clamp micropipette (see **Viab. Mecha. Death** (control 2)) led to a statistically significant 68% difference with FUS and Sham (*p <* 0.0001) (Fig. 2-h). These results indicate that FUS neurostimulation parameters capable of inducing *focal and propagating* Ca^2+^ responses did not compromise the immediate survival of targeted cells.

To understand the biophysical mechanisms behind the observed ultrasound-evoked Ca^2+^ uptake, *immediate* focal effects (≤ 1 s after FUS initiation) are reported first, followed by *delayed* propagating responses.

### 2.2 Influence of FUS frequencies on the focality of the immediate Ca^2+^ responses

The feasibility of evoking causal, focal and propagating Ca^2+^ responses with 400 µs single-pulse FUS protocols was tested throughout a wide range FUS frequencies (from 0.73 to 8.14 MHz) with a unique transducer driven at its base frequency and harmonics (Fig. 3-a-b). Using single-pulse FUS at *f*_0_ = 0.73 MHz led immediately after stimulation to the initiation of Ca^2+^ activity in multiple cell clusters (see **FUS 0.7** group) (Fig. 3-e). At this frequency, the diameter of the − 3 dB FUS focal spot spanned 2.5 mm (Fig. 3-c). Cluster diameters and inter-cluster distances were measured at 120 ± 20 µm and 940 ± 590 µm respectively (Fig. 3-e-g). Although their centroids were contained within the FUS focal spot, their locations and dimensions were random and did not clearly matched the focal spot spatial distribution. These random distributions were also found on the qualitative displacement maps reconstructed experimentally after single-pulse FUS emissions on optical phantoms (Fig. 3-d). This implied that the multi-cluster Ca^2+^ patterns observed on the neural cells were not only a consequence of intrinsic biological mechanisms, but also conditioned by FUS physical interactions with the neural cell network. Along with these observations, high inertial cavitation indices (30 ±1 dB) were measured during the FUS stimulation (Fig. 3-h). Low thermal elevations (spatial and temporal peak value) were estimated via numerical modeling, both in the Polydimethylsiloxane (PDMS) subtract of the cells (Δ*T*≪ 0.1^°^C) and in the cell layer (Δ*T* ≈ 0.0^°^C) (Table 1). Increasing the frequency of single-pulse FUS to *f*_3_ = 2.23 MHz with the same transducer allowed eliciting similar activity patterns with comparable cluster diameters of 100 ± 50 µm (see **FUS 2.2** group). At this frequency, the diameter of the focal spot decreased to 710 µm. Inter-cluster distances halved to 500 ± 200 µm, again corresponding to the emergence of several clusters in the focal spot. Additionally, inertial cavitation indices dropped by 13% and the PDMS substrate reached a peak temperature elevation of Δ*T* = 2.2 ± 2.0^°^C, which resulted in Δ*T* = 0.5 ± 0.5^°^C elevations in the cell layer (*Suppl. Video FUS 2*.*2*).

**Table 1.** Acoustic and thermal quantities obtained via numerical simulations based on acoustic powers allowing the initiation of propagated Ca^2+^ responses across the range of tested ultrasound central frequencies.

| FUS frequencies [MHz] | 0.73 | 2.23 | 5.19 | 8.14 |
| --- | --- | --- | --- | --- |
| −3 dB focal spot lateral diameter [mm] | 2.52 | 0.71 | 0.29 | 0.19 |
| −6 dB focal spot lateral diameter [mm] | 3.31 | 1.03 | 0.39 | 0.26 |
| $p_{\text{spr}}$ [MPa] | $0.78 \pm 0.31$ | $7.78 \pm 3.64$ | $8.79 \pm 2.10$ | $9.30 \pm 1.30$ |
| $p_{\text{sar}}$ [MPa] | $0.66 \pm 0.27$ | $6.59 \pm 3.08$ | $7.52 \pm 1.80$ | $8.01 \pm 1.12$ |
| $I_{\text{sppa}}$ [$\text{kW.cm}^{-2}$] | $0.18 \pm 0.11$ | $4.21 \pm 3.91$ | $6.89 \pm 3.26$ | $7.17 \pm 2.08$ |
| $I_{\text{sapa}}$ [$\text{kW.cm}^{-2}$] | $0.14 \pm 0.08$ | $2.93 \pm 2.72$ | $5.22 \pm 2.47$ | $5.45 \pm 1.58$ |
| $E_{\text{fspot}}$ [mJ] | $2.75 \pm 1.68$ | $5.09 \pm 4.73$ | $1.31 \pm 0.62$ | $0.55 \pm 0.16$ |
| $E_{\text{tot}}$ [mJ] | $14.39 \pm 8.80$ | $8.57 \pm 7.96$ | $3.91 \pm 1.85$ | $1.68 \pm 0.49$ |
| $\Delta T_{\text{PDMS sptp}}$ [$^{\circ}\text{C}$] | $0.01 \pm 0.00$ | $2.50 \pm 2.32$ | $9.22 \pm 4.36$ | $18.45 \pm 5.36$ |
| $\Delta T_{\text{PDMS satp}}$ [$^{\circ}\text{C}$] | $0.01 \pm 0.00$ | $2.15 \pm 2.00$ | $7.73 \pm 3.66$ | $15.55 \pm 4.52$ |
| $\Delta t_{\Delta T \text{ cells sptp}}$ [$\mu\text{s}$] | — | $399.55 \pm 0.00$ | $399.81 \pm 0.00$ | $399.88 \pm 0.00$ |
| $\Delta T_{\text{cells sptp}}$ [$^{\circ}\text{C}$] | $0.00 \pm 0.00$ | $0.61 \pm 0.57$ | $3.61 \pm 1.71$ | $7.39 \pm 2.15$ |
| $\Delta T_{\text{cells satp}}$ [$^{\circ}\text{C}$] | $0.00 \pm 0.00$ | $0.52 \pm 0.48$ | $3.14 \pm 1.49$ | $6.42 \pm 1.87$ |
| $\Delta t_{\Delta T \text{ cells sptp}}$ [ms] | — | $17.36 \pm 4.72$ | $23.22 \pm 1.58$ | $7.56 \pm 0.11$ |
| $\frac{\Delta T_{\text{cells sptp}}}{\Delta t_{\Delta T \text{ cells sptp}}}$ [$^{\circ}\text{C.ms}^{-1}$] | — | $0.03 \pm 0.12$ | $0.15 \pm 1.08$ | $0.98 \pm 19.54$ |

**Figure 3.**
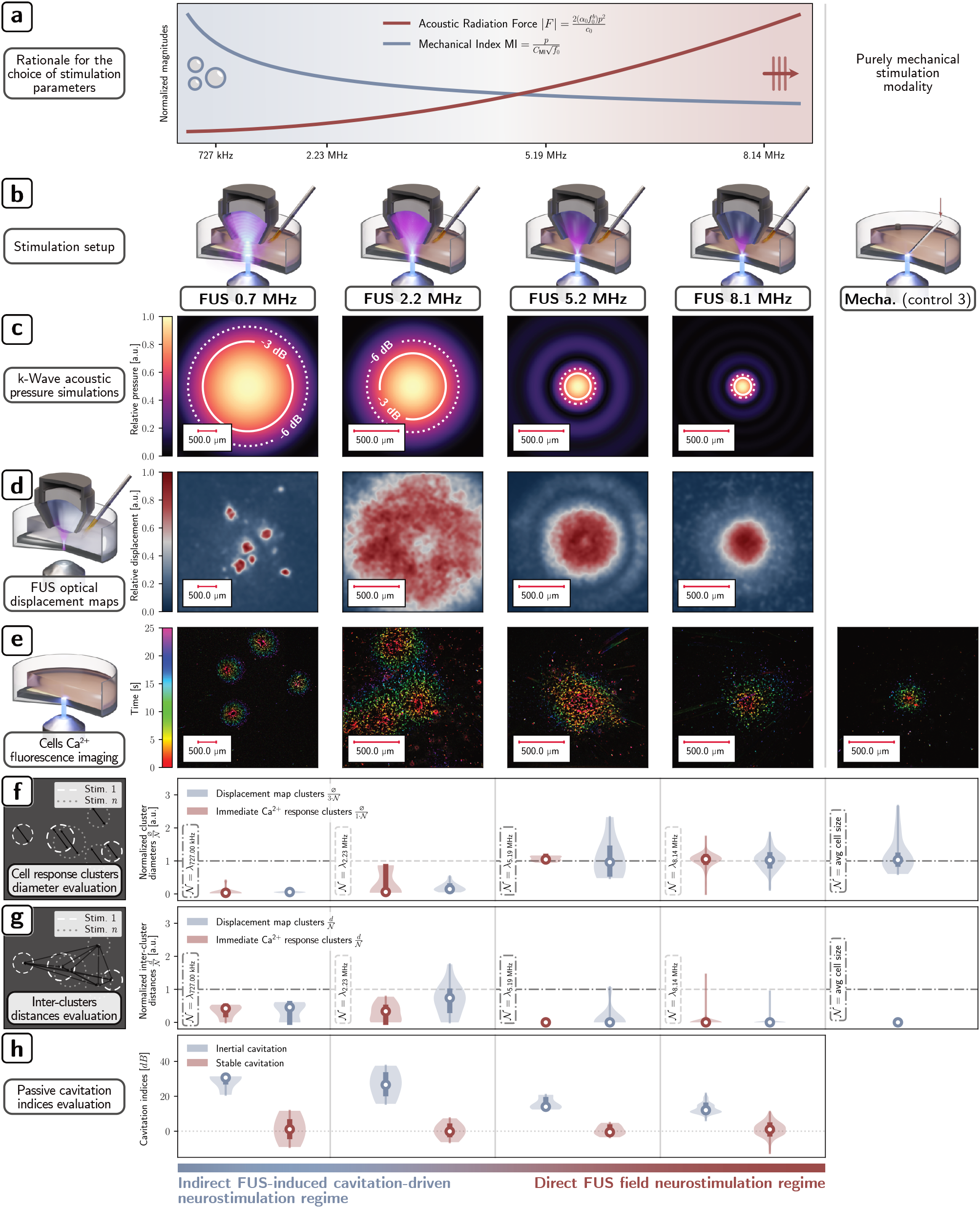
Influence of FUS frequencies on spatial selectivity and targeting accuracy in neurostimulation of immediate Ca^2+^ responses. **(a)** Qualitative illustration of the frequency dependence of the acoustic radiation force (ARF) |F| directly induced by the FUS field versus that of the mechanical index MI (indicative of the likelihood of triggering destructive effects such as cavitation events). This being the rationale for studying FUS neurostimulation in the 0.73-8.14 MHz frequency range. **(b-h)** From left to right, the columns respectively represent the type of data analysis performed followed by results obtained at 0.73 MHz (**FUS 0.7** group), 2.23 MHz (**FUS 2.2**), 5.19 MHz (**FUS 5.2**), 8.14 MHz (**FUS 8.1**) and with single-cell quasi-static mechanical stimulations (**Mecha**. (control 3) group). **(b)** Illustrations of the in vitro neurostimulation set-up including with 3D renders of the numerically simulated ultrasound fields in the case of FUS neurostimulation (from 0.73 to 8.14 MHz) or an illustration of a patch clamp micropipette in the case of single-cell quasi-static mechanical stimulation. **(c)** Spatial distribution of numerically simulated ultrasound pressure imposed by single-pulse FUS stimulation. **(d)** Spatial distribution of FUS-induced mechanical stresses evaluated qualitatively on an optical phantom under microscopy imaging and using optical particle tracking methods. **(e)** Biological responses of neural cells resulting from FUS and single-cell mechanical stimulations. Neural cell activity was assessed using Ca^2+^ fluorescence imaging and colored compound images were formed according to the process described in Fig. 2-f-g. Measurements characterizing the spatial distribution of immediate Ca^2+^ responses (displayed in red) and the spatial distribution of immediate mechanical stresses (displayed in blue) are reported. Measurements of Ca^2+^ responses were assessed from clusters of responding cells corresponding to the groups of cells colored red in the biological responses shown in (e). Mechanical stress measurements were assessed from the displacement maps in (d). **(f)** Normalized cluster diameters ϕ/ 𝒩 . **(g)** Normalized inter-cluster distances d/𝒩 reflecting the disperse nature of cavitation-elicited mechanical stresses and neural responses. **(h)** Stable and inertial cavitation indices reported for single-pulse FUS stimulations.

From *f*_7_ = 5.19 MHz, immediate responses were confined in a single cluster of responding cells, always centered on the FUS focal spot. At this frequency, the diameter of the focal spot could be refined to 290 µm. Cluster diameters were measured at 280 ± 110 µm, closely matching the focal spot diameter. Moreover, inertial cavitation indices declined to 14±2 dB while thermal elevation reached a peak of 9.2 ± 4.4^°^C in PDMS and 3.6 ± 1.7^°^C in the cells. Finally, among the tested frequencies, maximum neurostimulation spatial selectivity was reached at *f*_1_1 = 8.14 MHz (see **FUS 8.1** group). The diameter of the focal spot could be refined to 190 µm which also matched the diameter of the responding cells clusters (190 ± 30 µm). Temperature elevations in PDMS and cells peaked at Δ*T* = 18.4 ± 5.4^°^C and Δ*T* = 7.4 ± 2.1^°^C respectively. Interestingly, quasistatic single-cell mechanical stimulations, performed with micrometer-scale patch-clamp micropipette tips, elicited sustained Ca^2+^ activities equivalent to high frequency FUS stimulations occurring in single clusters 13 ± 1 µm in diameter, the size of a single cell body (Fig. 3, Table 1 and *Suppl. Videos FUS 5*.*2 & FUS 8*.*1*).

### 2.3 Two competing mechanisms: FUS field radiation force vs FUS-induced cavitation

Normalized cluster diameters (*ϕ* /𝒩) and intercluster distances (*d*/𝒩) of the immediate responses revealed the existence, within the *propagating* regime, of two distinct - *focal* or *dispersed* - frequency-dependent sub-regimes of Ca^2+^ activity initiation (being the characteristic dimension of the stimulus: the wavelength *λ* for **FUS** groups or the body cell size for the **Mecha**. (control 3) group). On the one hand, low-frequency FUS stimulations (≤ 2.23 MHz) resulted in multiple clusters of Ca^2+^ activity, randomly dispersed within the focal spot and associated to similar normalized distances (median *d*/ 𝒩 ≈ 1/2), corresponding to cluster distances growing, from 0.73 to 2.23 MHz, with the wavelength (median *d* ≈ *λ*/2) (**FUS 0.7 & FUS 2.2** groups, Fig. 3-g). In this frequency range, normalized cluster diameters were significantly smaller than 1 (median /𝒩 ≪ 1) which corresponds to clusters significantly smaller than the wavelength (median *ϕ* ≪ *λ*) (Fig. 3-f). On the other hand, high-frequency FUS (≥5.19 MHz) were similar to quasi-static mechanical stimulations (**FUS 5.2, FUS 8.1 & Mecha**. (control 3) groups). They both resulted in a single cluster of Ca^2+^ activity, centered within the focal spot (median *d*/𝒩 ≈ 0) with reduced variability in its localization throughout trials. The normalized cluster diameters were close to 1 (median /𝒩 ≈ 1) which translates to cluster diameters equivalent to the wavelength for FUS (median *ϕ* ≈ *λ*) or to the cell size for quasi-static mechanical stimuli (median *ϕ* ≈ cell diameter). The limit case of a cluster composed of a single cell was thus reached with quasi-static mechanical stimulation (Fig. 3-f).

These observations suggested that for *f <?*2.23 MHz, dispersed and random Ca^2+^ neurostimulations were evoked by indirect effects of the FUS field via stochastic cavitation. Indeed, no correlations were observed between: *1*. the controlled spatial distributions of the FUS pressure field and *2*. the uncontrolled dispersed spatial distributions of FUS-induced tissue deformations and Ca^2+^ responses (Fig. 3-c-e). In these conditions, stochastic cavitation effects appeared to prevail on controllable direct effects of the FUS field (Fig. 3-a). This was supported by passive cavitation indices, representative of the probability of presence of stable and inertial cavitation. Neither a significant presence nor a significant variation in the presence of stable cavitation was detected over the frequency range investigated. However, an inverse proportional relationship was found between *f* and inertial cavitation indices. In addition to the uncontrollable and dispersed nature of the Ca^2+^ responses, meaningful insights on the biophysical mechanisms - mechanical or/and thermal - associated to these stochastic inertial cavitation events can not be obtained from the linear acoustic modeling conducted in this study. Conversely, all observations made for *f* ≥ 5.19 MHz, pointed to direct a neurostimulation effect of the FUS field, and no significant role for cavitation. Single focal and immediate Ca^2+^ responses were repeatably evoked, with spatial locations and distributions well correlated to the imposed FUS pressure fields and estimated FUS-induced tissue deformations (Fig. 3-c-e). These observations - *a fortiori* correlated to results obtained with purely mechanical stimulations - suggested a predominant role of ultrasound mechanical effects in neurostimulation through FUS-induced radiation forces (Fig. 3-a). Interestingly, unlike Ca^2+^ responses observed on cell cultures, FUS-induced tissue deformations obtained in phantoms at 2.23 MHz showed spatial distributions coherent within a ratio of 3 with the FUS focal spot. This suggested that at this frequency, direct FUS effects could coexist with cavitation effects, can be predominant in some media containing few cavitation nucleis (e.g. optical phantom used here), or be surpassed by cavitation-driven effects in more heterogeneous acoustic domains (e.g. cell culture).

Overall, acoustic energies deposited within the FUS focal spot *E*_fspot_ and capable of evoking Ca^2+^ responses remained in the order of a mJ. The wide variations in the FUS spatial distributions and the high frequency-dependence of material properties (e.g. acoustic attenuation) used as a cell substrate (PDMS, glass petri dishes) resulted in a broad range of peak pressures, intensities, and associated temperature increases which were summarized in Table

1. Interestingly, neurostimulation directly evoked by the FUS field (*f*≥ 5.19 MHz) were associated to similar FUS intensity and pressure thresholds (*I*_sapa_ ≈ 5 kW.cm^−2^, *p*_sar_ ≈ 7 − 8 MPa). When cavitation was predominant (*f* = 0.73 MHz), the intensity and pressure levels estimated numerically for the FUS field only (no cavitation accounted in the model) were significantly lower by a ratio ≈ 35 −40 for intensities and ≈ 10 for pressures (*I*_sapa_ ≈ 0.14 kW.cm^−2^, *p*_sar_ ≈ 0.7 MPa). This also supported that direct effects of the FUS field were unlikely to be predominant at this frequency. At 2.23 MHz, where direct and indirect effects of FUS coexisted with variable predominancies, FUS pressure levels were similar in neural cells to those associated to successful neurostimulation (*psar*≈ 6 MPa), while FUS intensities where inferior by a ratio 2 (*I*_sapa_ ≈ 2 kW.cm^−2^). Since ultrasound intensity is proportional to the radiation force and the negative pressure is directly related to the initiation of cavitation, this was also supporting the predominance of cavitation in inducing neurostimulation at this mid frequency. Although all above observations converged on predominant mechanical effects of FUS, the hypothesis of a FUS-induced thermal effect was also considered. FUS-induced heating remained limited to the highly attenuating PDMS layer used as a mechanically compliant cell substrate and no correlations were observed between the various FUS-induced heating (peaks Δ*T*_cells sptp_ = 0 − 7^°^C for *f* = 0.73 − 8.14 MHz) and the minimal thresholds of neurostimulation (Table 1). All observations detailed in supplementary materials (Suppl. Results 1) were not in favor of an intrinsic effect of FUS-induced heating in the observed single-pulse FUS neurostimulation phenomena.

### 2.4 Neurobiological pathways involved in *focal* and *propagating* Ca^2+^ responses

We then sought to get insights on the neurobiological mechanisms mobilized by *f* = 8.14 MHz FUS stimulations, the latter being the most selective and repeatable FUS neurostimulation configuration studied. Following the immediate focal response described in the previous section, a slow, *delayed* intercellular Ca^2+^ flux propagation was observed in the surrounding region. In these trials, the assumption of a steady cell culture media (**no flow** condition) was made as the artificial Cerebro-Spinal Fluid (aCSF) perfusion was turned of. Both 8.14 MHz FUS and quasi-static mechanical stimulations evoked omnidirectional intercellular neural responses propagating at velocities *v*_no flow_(*t*) ranging from 20 to 30 µm.s^−1^ in the first five seconds ([1 − 5] s) before slowing down rapidly at ≈ 5 µm.s^−1^ after 25 s (**Flow off** (control 4) & **Mecha**. (control 3) groups) (Fig. 4). From the physics perspective, dynamics of the delayed propagating Ca^2+^ response revealed a 2D diffusion-like process with a velocity following a 1/*r* law (where *r* corresponds to the radius of the intercellular response). This 2D diffusion-like dynamic was modeled as a Bernoulli random walk [38] as a function of *ϕ*, the diameter of the immediate focal response, and a constant surface expansion rate *e* such that

**Figure 4.**
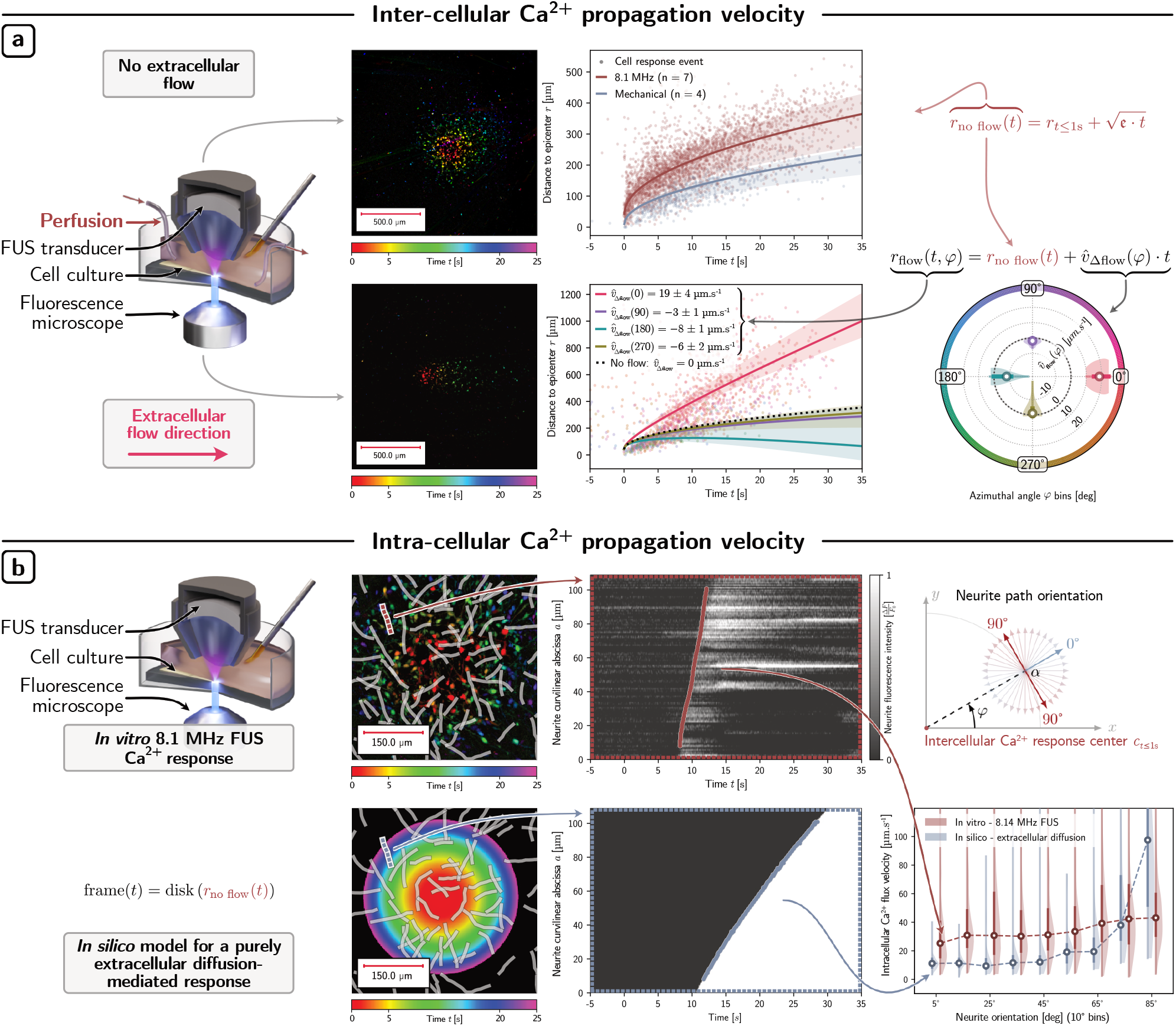
Spatiotemporal dynamics of FUS-induced Ca^2+^ responses and insights into the extra- and intra-cellular neurobiological pathways involved. **(a-b)** From left to right, columns represent respectively the experimental set-ups established to assess inter- and intra-cellular Ca^2+^ propagation velocities after FUS stimulations, the colored compound images of typical responses obtained, the representations of the spatio-temporal dynamics of the evoked Ca^2+^ responses, radial distances as a function of time in section (a) (intercellular dynamics) or curvilinear propagation distances as a function of time in section (b) (intracellular dynamics). Propagation velocities as a function of the propagation direction in the FOV are then shown in section (a) (intercellular dynamics) followed by intra-neurite Ca^2+^ wave propagation velocities as a function of their orientation relative to the center of the intercellular response (intracellular dynamics). **(a)** Intercellular dynamics of 8.14 MHz FUS-induced (in red) vs mechanically-induced (in blue) Ca^2+^ responses obtained in static cell culture medium (top row, **Flow off** (control 4) vs. **Mecha.** (control 3) groups) and after imposing a lateral extracellular flow to the medium (bottom row, **Flow on** (control 5)). An isotropic form of an expansion model was fitted to responses evoked by the FUS without any flow [38]. While dots correspond to response events of individual cells in the FOVs, fit results are represented by solid lines on the graphs. The Q_1_ − Q_3_ interquartile ranges are represented by the filled-in areas. Individual cell responses in the presence of lateral flow were grouped into bins of 0^°^, 90°, 180° and 270° centered on φ and analyzed by fitting the anisotropic term 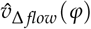 added to the result obtained without any flow. **(b)** Intracellular dynamics of FUS-induced Ca^2+^ responses at 8.14 MHz obtained in static cell culture medium. Experimental results of FUS-induced Ca^2+^ responses overlayed with hand-drawn neurite paths along which intracellular Ca^2+^ curvilinear velocities were evaluated (top row, **Flow off** (control 4) group). These were compared with an in silico model assuming purely extracellular signaling pathway (bottom row). Finally, the distribution of intracellular Ca^2+^ flux velocities was reported for a range of neurite path orientations, confirming that experimental observations diverged from the hypothesis of a pure extracellular pathway.

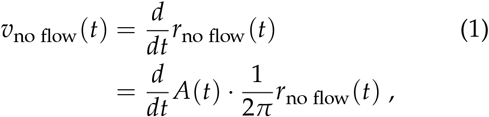

where 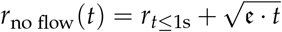 with 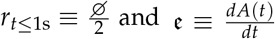 a constant. *r*_no flow_(*t*) corresponds to the expansion radii of Ca^2+^ responses without flow, *A*(*t*) to the cumulative response area covered by all neural cells having responded to neurostimulation at a time *t* and 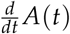 the response area expansion rate (Fig. 4-a and *Suppl. Videos Flow off (control 4) & Mecha. (control 3)*).

From the biological perspective, this diffusion-like dynamic could be attributed to both intra- and extra-cellular signaling pathways. A first limit case of an *exclusively extracellular diffusion* was considered to consist in the extracellular release of chemical messengers within the focal response cell cluster during the immediate phase (*t* ≤1 s) of the response, triggered new delayed Ca^2+^ cell activity as the chemical messenger diffuses in the extracellular media. On the other hand, a *purely intracellular propagation* was assumed to correspond to a signaling mechanism taking place within the cells themselves. In the hypothesis of a purely intracellular pathway, sampling Ca^2+^ flux velocities along elongated the trajectory of cells bodies is expected to lead to velocities largely independent from the propagation direction, solely related to the conduction velocity of cells. In this limit case, the diffusion-like dynamic of the intercellular propagation would arise from the convoluted architecture of the cellular network. From a physics perspective, this corresponds to a random walk process occurring at the scale of the cellular network. In contrast, the property of the velocity being independent from the trajectory direction no longer holds when the same reasoning is applied to the case of purely extracellular signaling. In deed, when considering the velocity evaluated on a path parallel to the intercellular Ca^2+^ wavefront, it is clear that fluorescence elevation would occur simultaneously along the path, leading to an apparent velocity approaching infinity. In the frame of the random walk model, this emergent property can be justified when dealing with a process taking place on a much smaller scale than that of the observer. Indeed, the diffusion of an extracellular messenger can be seen as a random walk process occurring on a molecular scale.

To discriminate the scale of the processes at play in FUS-evoked propagating waves, Ca^2+^ fluxes velocities were thus evaluated along identifiable cell neurites after FUS stimulation. Near-constant intracellular velocities of 30 ± 15 µm.s^−1^ were measured and no clear relationship was found between these velocities and the orientation of cell bodies and neurites (Fig. 4-b). Performing the same measurement on the *in silico* model of a purely extracellular diffusion led to results notably different from experimental analyses. In this limit case, velocities sampled along paths appeared to strongly dependent on their orientations, with values increasing significantly as the paths orientations approached 90^°^ (Fig. 4-b). This result rejected the purely extracellular hypothesis, and supported the intracellular pathway hypothesis in which a network-scale random-walk process could be at the root of the diffusion-like propagation dynamic.

However, the addition in some experiments of a forced extracellular flow using a perfusion pump had significant effects on the overall Ca^2+^ wave propagation (**Flow on** (control 5) group) (Fig. 4-a). Introducing the parameter *φ* ∈ [0, 360]^°^ in the response expansion model (equation 2) allowed to describe the propagation anisotropy. In this configuration, intercellular Ca^2+^ propagation velocities increased by 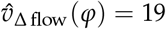 µm.s^−1^ or slowed down by 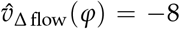 µm.s^−1^ in the direction of flow (0^°^ orientation) and its opposite (180^°^ orientation) respectively. Notably, orthogonal directions were comparatively unaffected. Experimental results obtained in these experimental conditions (**flow**) were successfully fitted in the expansion model by introducing the *φ*-dependent term 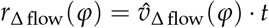, defining the variation of intercellular Ca^2+^ expansion radius as a function of the drift of Ca^2+^ expansion velocity induced by the perfusion such that

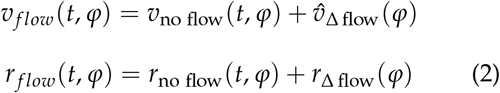

where *r* _*f low*_(*t, φ*) and *v* _*f low*_(*t, φ*) respectively the expansion radius and velocities of the responses in presence of an extracellular flow. These results suggested the coexistence of two distinct intra- and extra-cellular signaling pathways in the intercellular Ca^2+^ flux propagation induced by 8.14 MHz single-pulse FUS neurostimulation (Fig. 4-a and *Suppl. Videos Flow off (control 4) & Flow on (control 5)*).

### 2.5 Cellular mechanisms regulating FUS-evoked Ca^2+^ responses

Pharmacological blockade experiments allowed the identification of cellular mechanisms regulating the neurobiological signaling pathways involved in 8 MHz single-pulse FUS-evoked intercellular focal and propagating Ca^2+^ responses (Fig. 5). The role of action potentials (APs) in the signaling process was first investigated by introducing the Na^+^ channels blocker TTX, resulting in a non-statistically significant 34% reduction of individual cell fluorescence elevation rates compared to control conditions (**TTX + flow off** group; Fig. 5-b, 1^st^ row). The neurostimulation success rates (*NSR* = 78%) remained stable compared to control (**Flow off** (control 4) group), with a limited reduction of the percentage of cells immediately responding in the focal spot (Fig. 5-c). Overall, the propagation of Ca^2+^ responses was unaffected, with no notable difference in the area expansion ratio 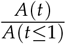 of propagating responses (Fig. 5-b, 2^nd^ row). These data suggested that neuronal APs were not required to evoke both *immediate focal* and *delayed propagating* Ca^2+^ responses. Given previous descriptions mechanically-induced Ca^2+^ responses [35, 39, 40], implication of a glutamate-mediated extracellular signaling pathway was investigated by introducing the NMDA receptor antagonist d-AP5 (**d-AP5 + flow** group). This resulted in a significant slowdown of the fluorescence elevation rates for cells located in the focal spot (97% reduction of the peak cell fluorescence elevation rate, max 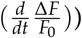. Responding clusters remained however detectable with a NSR of 89%. Most notably, the sensitivity of the propagation to an extracellular flow vanished in the **d-AP5 + flow** group. Overall, propagating responses were also significantly altered with a 95% reduction of the area expansion ratios, suggesting that glutamate is one of the primary messengers involved in 8 MHz single-pulse FUS-evoked extracellular Ca^2+^ response diffusion (*Suppl. Videos TTX + flow off, d-AP5 + flow & Flow off (control 4)*).

**Figure 5.**
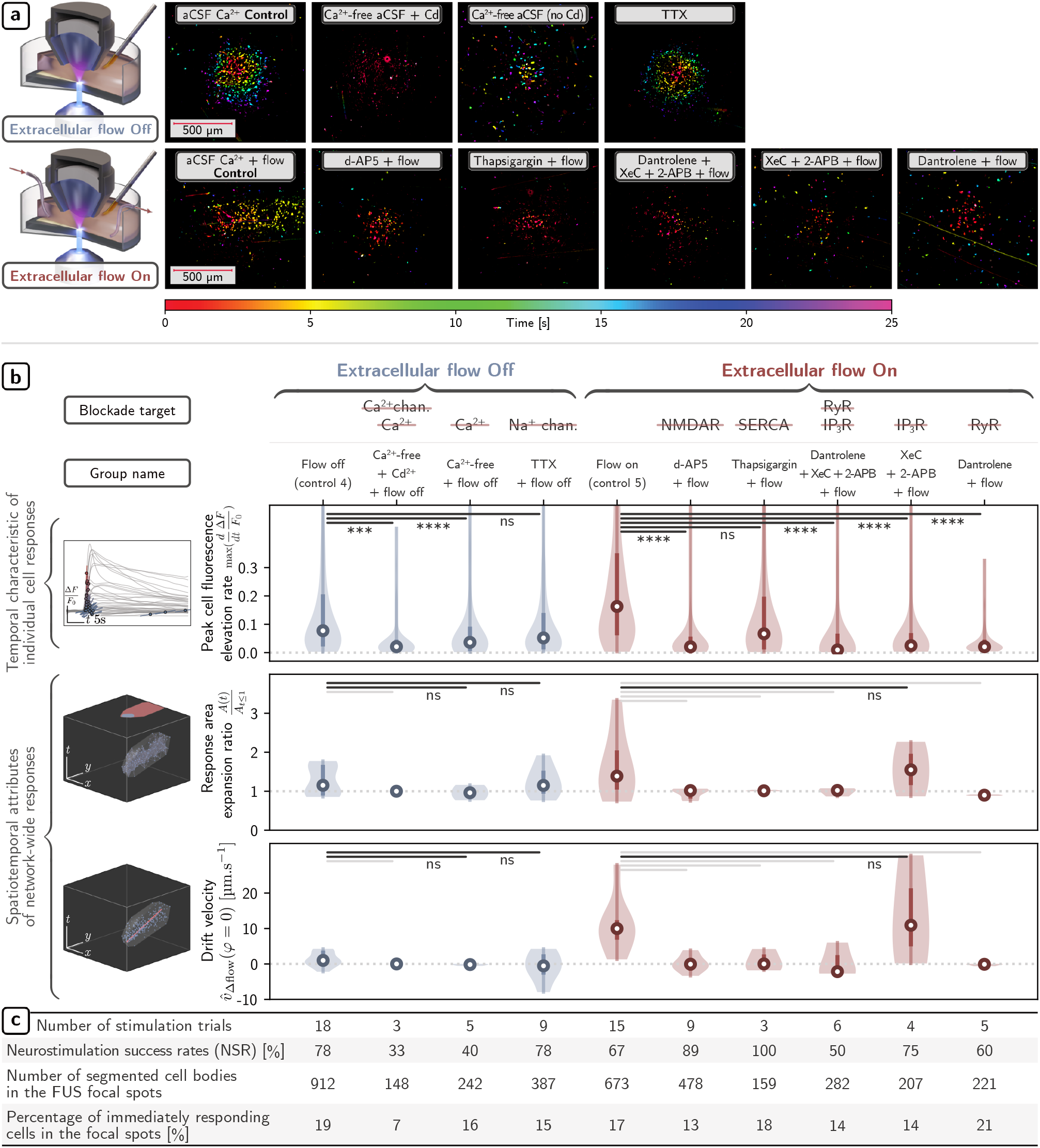
Cellular mechanisms regulating FUS-induced Ca^2+^ responses and their intercellular propagation involving both intra- and extra-cellular pathways. (a) Experimental set-ups and colored compound images representing the results of typical Ca^2+^ responses obtained after 400 µs single-pulse 8.14 MHz FUS neurostimulation under various pharmacological blocking conditions, with and without extracellular flow. On the upper row, FUS stimulations without extracellular flow and from left to right respectively: without blockade (see Flow off (control 4) group), in the absence of extracellular Ca^2+^ and inactivating Ca^2+^ channels (Ca^2+^-free + Cd^2+^ + flow off), in the absence of extracellular Ca^2+^ but with active Ca^2+^ channels (Ca^2+^-free + flow off), or in the presence of extracellular Ca^2+^ while blocking AP creation by inactivating Na^+^ channels (TTX + flow off). On the bottom row, FUS stimulation with additional extracellular flow and from left to right respectively: without blockade (see Flow on (control 5) group), with an NMDA receptor antagonist to block any implication of an extracellular glutamate messenger (d-AP5 + flow), while inhibiting ER SERCA pumps (Thapsigargin + flow), while blocking upstream processes regulating Ca^2+^ release in the ER with inactivation of IP_3_R and Ryanodyne receptors (RyR) (Dantrolene + XeC + 2-APB + flow), while blocking IP_3_R receptors alone (XeC + 2-APB + flow), or while blocking RyR alone (Dantrolene + flow). (b) Statistical analyses of FUS-induced Ca^2+^ responses obtained under different pharmacological blockade conditions, with and without extracellular flow. From left to right, the type of data analysis and plots pooling the results for each group are shown. Three parameters are evaluated: the peak cell fluorescence elevation rate for individual cell bodies located in the FUS focal spot (top row), the response area expansion ratio (middle row) and the response velocity drift along the angular direction (φ = 0^°^) of the extracellular flow (bottom row). Propagating Ca^2+^ responses correspond to expansion ratios >1. Assuming isotropic cell architecture, non-zero drift velocity medians were interpreted as the susceptibility of a response to extracellular flow. (c) Summary for each pharmacological blockade condition - with and without extracellular flow - of the number of stimulation trials, NSRs, number of cell bodies segmented in the FUS focal zone and percentage of cells responding immediately to FUS stimulation in the focal zone.

The implication of intra-cellular Ca^2+^ stores, namely the endoplasmic reticulum (ER), was then investigated. Since massive Ca^2+^ release from intracellular ER stores have already been reported to elicit glutamate release through exocytosis or dedicated channels like BEST-1 [35], theses mechanisms were tested in the context of single-pulse FUS neurostimulation. Intracellular ER-Ca^2+^ stores blockade was conducted at three levels: *1*. through ER-Ca^2+^ store depletion by blocking Ca^2+^ uptakes within ER via the inhibition of SERCA pumps located on the ER endomembrane using Thapsigargin or *2*. by blocking upstream processes regulating the Ca^2+^ ER release via the blockade of intracellular Ca^2+^ channels of the ER endomembrane. Either *2a*. Inositol 1.4.5-triphosphate receptors (IP^3^R) with XeC + 2-APB and/or *2b*. Ryanodine receptors (RyR) with Dantrolene. The depletion of ER-Ca^2+^ stores led to a 35% slowdown in the fluorescence elevation rates, reduced responses expansion ratios by 27% and eliminated the susceptibility to an extracellular flow (**Thapsigargin + flow** vs **Flow on** (control 5) groups). These results confirmed the implication of ER-Ca^2+^ stores in the propagation of Ca^2+^ intercellular responses. On the other hand, the blockade of upstream processes regulating the Ca^2+^ ER release (IP^3^R and RyR) led to a major and statistically significant 98% slowdown of the fluorescence elevation rates vs Control 5 (*p <* 0.0001) (**Dantrolene + XeC + 2-APB + Flow** group; Fig. 5-b, 1^st^ row). Similar to what was observed when depleting ER-Ca^2+^ stores, blocking those two receptors involved in the Ca^2+^ ER release resulted in a 26% reduction of the area expansion ratios and eliminated the susceptibility to the extracellular flow (Fig. 5-b, 2^nd^ row). The blockade of IP_3_R alone, however, did not alter response propagation in the same way. Results indicated limiting effects (11% reduction) in the area expansion ratios and the susceptibility to the extracellular flow vs. Control 5 was maintained (only 9% reduction of the drift velocities) (**XeC + 2-APB + flow** group). On the other hand, the blockade of RyR alone (**Dantrolene + flow** group) suppressed all intercellular propagation and eliminated the susceptibility to the extracellular flow, which was comparable to what was observed when blocking both IP_3_R and RyR (Fig. 5-a). Overall, these results clearly indicated the implication of Ca^2+^ release from intracellular ER stores in FUS-evoked propagating responses, and more specifically, the role of RyR in the triggering of the release (*Suppl. Videos Thapsigargin + flow, Flow on (control 5), Dantrolene + XeC + 2-APB + Flow, XeC + 2-APB + flow & Dantrolene + flow*).

Finally, in order to characterize the mechanisms leading to RyR activation and subsequent Ca^2+^ release, the implication of calcium ion channels of the plasma membrane was investigated. The point was to determine whether FUS can evoke Ca^2+^ influx either *1*. by acting directly on the RyR Ca^2+^ channel within the ER endomembrane, or *2*. by indirectly acting on the functions of these RyRs by affecting the intracellular Ca^2+^ messenger via previously reported calcium-induced calcium release (CICR) amplification process involving extracellular Ca^2+^ entry in the cell through plasma Ca^2+^ channels [41, 42]. Single-pulse FUS stimulations were thus conducted using altered extracellular aCSF media. Replacing the extracellular Ca^2+^-rich aCSF medium with a Ca^2+^-free aCSF medium while adding the Ca^2+^ channel blocker cadmium Cd^2+^ had a drastic impact on the NSR, reducing it by more than half vs Control 4 (**Ca**^**2+**^**-free + Cd**^**2+**^ **+ flow off** group). Fluorescence elevation rates for cells located in the focal spots were also significantly slowed down by 92% and no response propagation could be measured afterwards (Fig. 5-a and 5-b-1^st^ row). These observations suggested that extracellular Ca^2+^ entry was required to initiate FUS-evoked intercellular Ca^2+^ signaling, by relying on RyR-mediated CICR amplification processes. Interestingly however, without blockade of Ca^2+^ channels with Cd^2+^, FUS could still elicit Ca^2+^ responses in Ca^2+^-free aCSFs (**Ca**^**2+**^**-free + flow off** group). Despite a significant 82% slowdown of fluorescence elevation rates and of response expansion ratios vs Control 4, this suggested that FUS gating of functional plasma Ca^2+^ channels allowed: *1*. either the re-entry of trace amounts of extracellular Ca^2+^ (released by cells after media replacement) or *2*. triggering another mechanism mediated by plasma Ca^2+^ channels and capable of gating RyR without extracellular Ca^2+^ entry. The initiation of large scale responses in Ca^2+^-free aCSFs provided further evidences for the involvement of Ca^2+^ amplification such as that being in play in CICR processes (*Suppl. Videos Ca*^*2+*^*-free + Cd*^*2+*^ *+ flow off & Ca*^*2+*^*-free + flow off*).

## 3 Discussion

### 3.1 First evidence of highly focal FUS neurostimulation on human neural cells

Following the definition of neurostimulation proposed by Gavrilov [6], the present study showed first evidence of highly selective FUS neurostimulation on human neural cells, with the causal initiation of *immediate focal* and *delayed propagating* Ca^2+^ neural responses. The study explored the effect of ultrasound frequencies, pressures, powers, or intensities with single, short FUS pulses (400 µs-long). By acquiring control over the initiation of causal neural activity in a spatially selective and safe way in human neural cells with elementary FUS stimuli, we expect to improve the therapeutic outcomes of complex repetitive-pulse neurostimulation sequences in the future. The presented context of transdural FUS intracranial brain stimulation implants for the management of chronic neurological diseases also allowed the investigation of a broad range of ultrasound frequencies, from sub-MHz, conventionally used in transcranial FUS (TUS) approaches, to much higher frequencies up to 8 MHz with a high potential to improve spatial selectivity.

### 3.2 Single-pulse FUS as an elementary contactless mechanical neurostimulation

Single-pulse FUS were test as an elementary brick of neurostimulation on a broad range of frequencies. Our approach allowed the unification of the two main hypotheses competing in the literature by validating two ultrasound effects able to induce neurostimulation: *1*. direct interaction of the FUS field with neural cells (*f* ≥ 5.19 MHz) and *2*. FUS-induced cavitation for *f* ≤ 2.23 MHz. The predominance of the two FUS neurostimulation regimes was shown to be frequency-dependent, which allowed the selection of FUS frequencies optimizing the efficacy, accuracy and safety the procedure. In the lowest frequency range (*f* ≤ 2.23 MHz), *dispersed* causal Ca^2+^ responses driven by inertial cavitation were randomly evoked within the FUS focal region, which raised several problems in terms of spatial targeting accuracy and safety. In the highest frequency range (*f* ≥ 5.19 MHz), Ca^2+^ activities were always directly initiated by the FUS field itself, within the FUS focal spot, and with negligible probabilities of cavitation. These configurations provided higher efficacy, accuracy and safety. The mechanical nature of the interactions responsible for the immediate responses evoked by single-pulse FUS was confirmed. A specific FUS neurostimulation regime was identified where repeatable mechanical interactions could be induced directly by the FUS radiation force (*f* ≥ 5.19 MHz). In this regime, *immediate focal* Ca^2+^ neural responses were characterized by maximal spatial selectivity and strong spatio-temporal causality. These immediate responses were followed by a *delayed propagating* intercellular Ca^2+^ wave. Importantly, no frequency effect has been noticed between 5.19 & 8.14 MHz with similar minimal neurostimulation thresholds, which was in favor of selecting the highest frequency, associated to the finest spatial selectivity (*<* 200 µm) and most robust strategy (78% NSR) while maintaining the neural cell viability. By promoting the action of the global radiation force at the focus, single-pulse megahertz-range FUS behaved as a contactless quasi-static mechanical stimulation. These findings allow considering the transdural FUS approach as a wearable mean of achieving deep brain mechanical stimulation without needle insertion in brain tissues.

### 3.3 Value of megahertz-range for transdural FUS neurostimulation approaches

For chronic neurological disorders which may require frequent (potentially daily) treatments, the TUS neuro-modulation/-stimulation approaches appear very challenging. Indeed, guidance via MRI or optical tracking currently used to ensure proper targeting with transcranial neurostimulation modalities (TMS, tDCS, TUS) both require a high degree of expertise and complex equipment. To overcome these obstacles, Carpentier *et al*. proposed in 2016 a small intracranial extradural implant to deliver unfocused ultrasound chronically in the context of blood brain barrier (BBB) opening [15]. In contrast to TUS, this wearable strategy coupled in our study to ultrasound focusing (FUS) opens the door to the use of higher frequencies (*>*5 MHz) associated, as we have shown, to acute neurostimulation spatial selectivity and reduction of cavitation-related adverse events. This approach has also been recently mentioned for sonogenetic neuromodulation of the visual cortex [16], which is promising for activating certain neural cell types selectively, although requiring genetic manipulation. In the present work, the single-pulse FUS strategy could allow the development of intracranial extradural FUS implants capable of activating clusters of cells with a spatial selectivity in the micro-metric scale, and without requiring genetic modifications. While encouraging, it is worth noting that our observations cannot be extrapolated directly to the 3D *in vivo* full brain, nevertheless, the identified trends on the ultrasound mechanism’s preponderance (cavitation vs FUS field effects), neural response thresholds and regimes (stable minimal FUS pressure/intensity thresholds at 5−8 MHz), optimization of the spatiotemporal focality and dynamics of the responses (focal and propagating) should carry. The possibility of increasing the frequency of short single-pulses FUS may also be of interest in terms of safety, as it will decrease the mechanical index 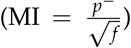. The stable minimum pressure thresholds observed between 5 and 8 MHz suggest that similar NSRs should be achievable by increasing frequencies while minimizing cavitation.

### 3.4 Direct action of FUS-induced radiation force evokes immediate focal Ca^2+^ responses

Most hypotheses formulated to elucidate the ability of ultrasound to induce Ca^2+^ influx in neural cells rely on the entry of extracellular Ca^2+^ into the cell’s cytosol via the plasma membrane [43]. As summarized in Fig. 6-I, associated FUS-mediated biophysical mechanisms were classified into two categories: *Ia*. direct mechanical opening of ion channels or the creation of pores by the mechanical action of FUS on the cell membrane (compressional or shear stresses) and *Ib*. indirect opening of voltage-dependent ion channels involving mechanoelectrical transduction mechanisms mediated by the mechanical action of FUS on the cell membrane. In the first category, several ion channels (e.g. TRPA1, TRPP1/2, TRPC1, Piezo1) classified as mechanosensitive have already been proposed to be involved in FUS neuro-modulation/stimulation [30, 35]. However, although single-pulse FUS are transient (a few hundred microseconds) and associated with low energy, exposure to high transient mechanical stresses could also be able to open ion channels that are not usually classified as mechanosensitive. Therefore, the hypothesis that FUS exposures are capable of opening selective or non-selective voltage-gated or ligand-gated channels is also viable. Another direct pathway for the entry of extracellular Ca^2+^ could be attributed to the opening of micropores by sonoporation resulting from violent cavitation events perforating the plasma membrane [44]. In the second category (indirect activations), and as summarized by Sassaroli *et al*. [45], several models for FUS-induced mechanoelectrical transductions have been proposed to explain neuro-modulation/stimulation, including intra-plasma-membrane cavitation activity (NICE Model, [26]), plasma membrane capacitance changes [27], piezo-or flexo-electric effects [46]. As shown by Plaksin *et al*. [25], thermal elevation of neural structures may also be responsible for these electro-acoustic transduction effects. However, in our study, neither neuronal APs nor cavitation activities were required in the neurostimulation regime associated with robust evocation of focal Ca^2+^ responses (*f* ≥ 5 MHz). Thus, the main hypothesis retained is the activation of mechanosensitive [30, 35] or stretch-activated [47, 48] Ca^2+^ ion channels by the purely mechanical action of FUS-induced radiation forces.

**Figure 6.**
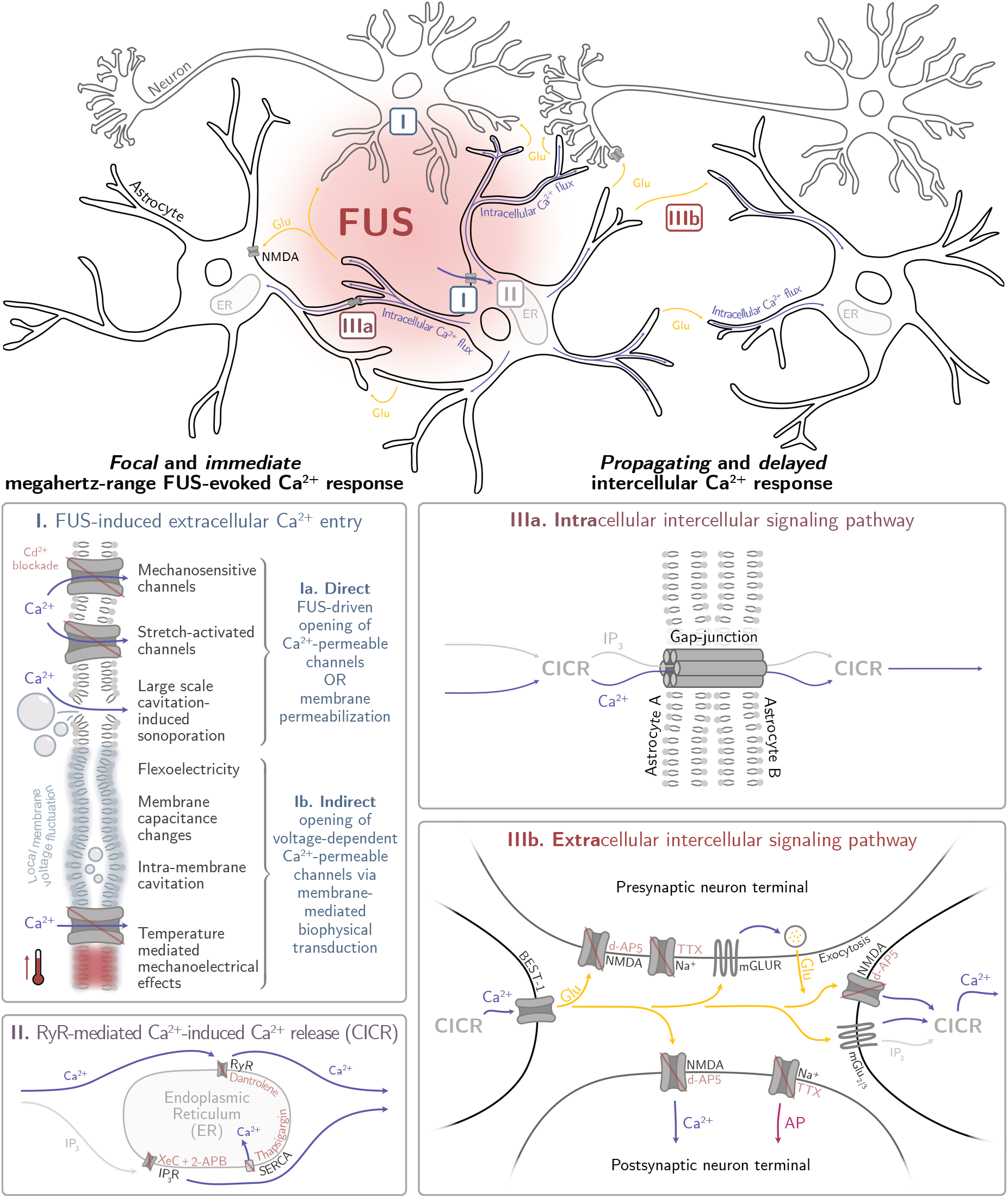
Current Hypotheses on Neurobiological Mechanisms underlying megahertz-range single-pulse FUS-induced immediate focal and delayed propagating Ca^2+^ responses. **(Top row)** Schematic representation of the whole chain of neurobiological mechanisms leading the the observed responses. This summary was established based on results obtained throughout this work by studying the spatiotemporal dynamic of slow Ca^2+^ and from the literature. **(I)** Gating of Ryanodine receptors (RyR) via direct or indirect FUS-induced Ca^2+^ entry through the plasma membrane. Although only limited data support this hypothesis for now, the existence of Ca^2+^-independent RyR activation mechanisms are discussed in Suppl. Discussion 2 (step **I bis**). **(II)** FUS-triggered RyR gating causes one or more mechanisms of intracellular amplification of RyR-mediated Ca^2+^ release, including Ca^2+^-induced Ca^2+^ release (CICR). **(III)** FUS-evoked CICR simultaneously drives **(IIIa)** an intracellular intercellular signaling pathway where Ca^2+^ diffusion can propagate within cells and from cell to cell via gap junctions and **(IIIb)** a Glutamate-mediated extracellular intercellular signaling pathway.

**Figure 7.**
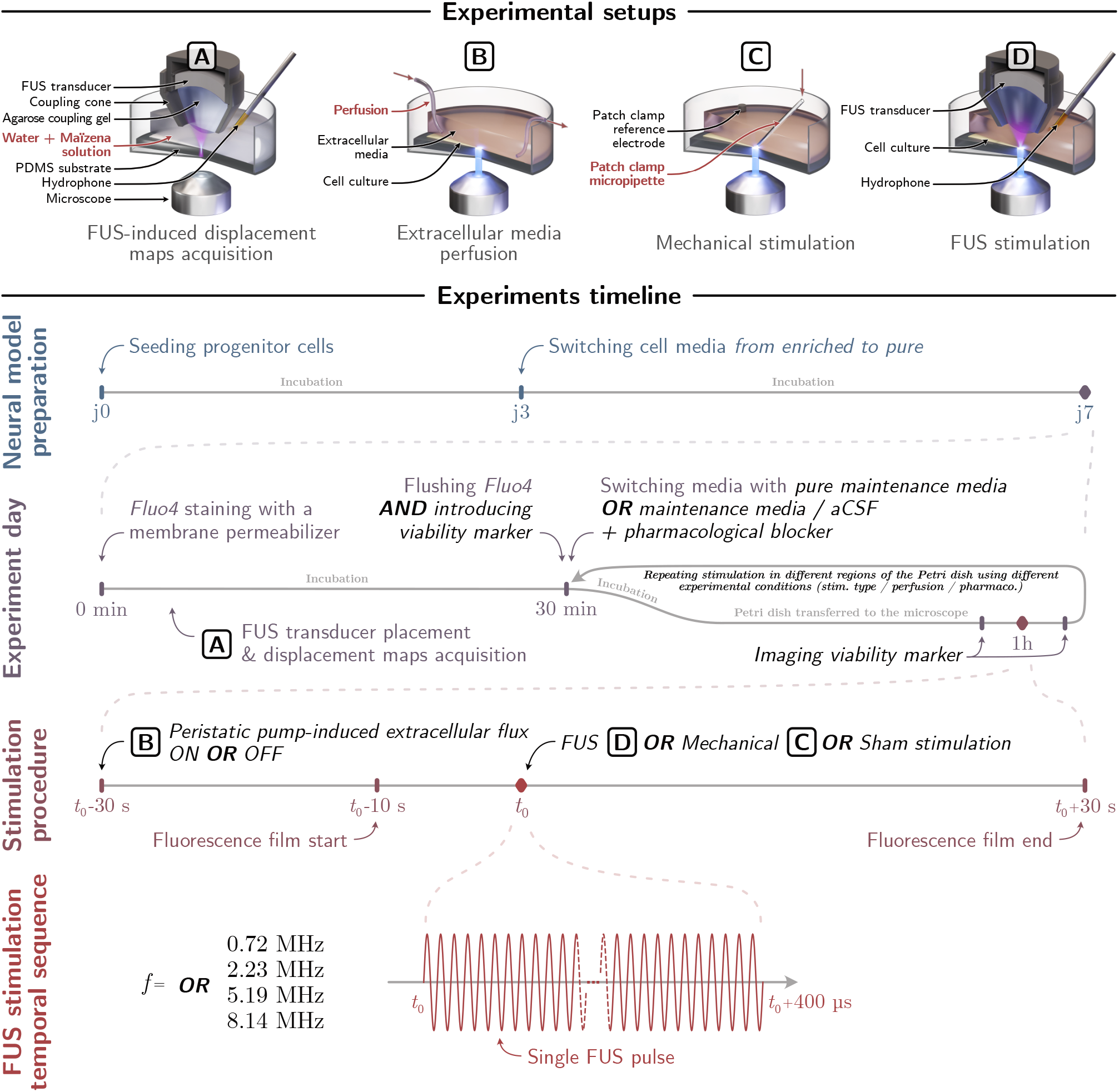
Overview of the experimental design and its temporal unfolding. (1^st^ row) Schematic view of the different in-vitro set-ups used in the presented study to evaluate respectively (A) the spatio-temporal distribution of single-pulse FUS-induced mechanical stresses on a tissue phantom; (B) the effect of additional extracellular flow on the spatio-temporal dynamics of FUS-induced Ca^2+^ responses; (C-D) the spatio-temporal dynamics of Ca^2+^ responses induced by (C) single-cell quasi-static mechanical stimulation and (D) single-pulse 400 µs FUS stimulation. (2^nd^ row) Chronology of the preparation of the human neural cell model in vitro; (3^rd^ row) Chronology on the day of the experiment to prepare cells and instruments to perform the neurostimulation protocols and neuromonitoring of induced Ca^2+^ waves; (4^th^ row) Chronology of the FUS neurostimulation protocol; (5^th^ row) the temporal sequences of single-pulse FUS stimulation used in the study at four different ultrasound frequencies.

### 3.5 Neurobiology of immediate focal and delayed propagating Ca^2+^ responses

We identified the following chain of neurobiological mechanisms which was consistent with standard descriptions of Ca^2+^ waves induced by quasi-static mechanical stimulations [49]. (Fig. 6). In step **II** (Fig. 6-II), FUS-evoked RyR gating was the main mechanism responsible for intracellular CICR amplification. CICR mechanism, originally described in cardiac muscle cells, was the hypothesis proposed by Yoo *et al*. [30] to explain FUS neurostimulation in cortical neurons. In cardiac muscle cells, CICR amplification is associated to: *1*. the opening of voltage-gated CaV1.2 plasma Ca^2+^ channel for *2*. the entry of extracellular Ca^2+^ in the cell leading to *3*. the gating of RyR2 isoform via a transient influx of extracellular Ca^2+^. This process was also described in neural cells to explain the initiation of intercellular Ca^2+^ wave propagation [49]. In our study, however, functional plasma Ca^2+^ channels were mandatory in FUS-evoked Ca^2+^ responses, while the role of extra-cellular Ca^2+^ re-entry into the cell was less clear. The observation of Ca^2+^ responses with Ca^2+^-free aCSFs could mean that remaining traces of extracellular Ca^2+^ could be sufficient to trigger CICR amplification. Thus, the involvement of RyR and the initiation of CICR by the entry of extracellular Ca^2+^ (even traces) via plasma ion channels opened by FUS is one main hypothesis (step **I**, Fig. 6-I). An alternative hypothesis where the initiation of the Ca^2+^ amplification does not require extracellular Ca^2+^ entry into the cell was also formulated in Suppl. Discussion 2 (step **I bis**).

In all the hypotheses, changes in intracellular Ca^2+^ concentration [Ca^2+^]i are likely to lead to subsequent CICR amplification processes (step **II**, Fig. 6-II). The temporal dynamics of the transient influx of Ca^2+^ and [Ca^2+^]i must, however, be controlled to avoid calcium-dependent inactivation of CICR processes [50], and the impact of FUS parameters on these mechanisms remains to be investigated. Finally, the intercellular propagating Ca^2+^ responses (step **III**, Fig. 6-III) involved the parallel implications of intra-and extra-cellular intercellular Ca^2+^ signaling pathways (Fig. 6-IIIa and 6-IIIb). Two messengers, Ca^2+^ and IP_3_, have been previously reported as individually capable of underlying the intracellular pathway via their diffusions through gap junctions [49]. Since FUS-evoked Ca^2+^ fluxes were found to be RyR-dependent and IP_3_R-independent, the Ca^2+^ messenger hypothesis was preferentially considered. It has also been reported that Ca^2+^-mediated Ca^2+^ fluxes can be distinguished from IP_3_-mediated ones on the basis of their much slower diffusion dynamics (in an oocyte model, ≈ 10 µm^2^.s^−1^ and ≈ 300 µm^2^.s^−1^ for Ca^2+^ and IP_3_ respectively) [49, 51]. In our study, intracellular Ca^2+^ propagation occurred at a constant velocity of ≈ 30 µm.s^−1^ inside neurites, which reinforced the hypothesis of a first intracellular intercellular pathway mediated by the Ca^2+^ messenger diffusing via gap junctions (Fig. 6-IIIa). To describe the second extracellular pathway, a mechanism is proposed for which the omnidirectional diffusion-like dynamics of responses emerged from the random and isotropic nature of the cellular network. As demonstrated by Charles *et al*. in astrocytes [40], intercellular Ca^2+^ signaling can also occur in disconnected cell cultures, revealing the existence of an extracellular messenger. Our observation that the responses was directly influenced by an extracellular flow led to the same conclusion for FUS-induced Ca^2+^ waves. This echoed observations made by several groups such as Innocenti *et al*. [39] on the involvement of glutamate, released by mechanically stimulating astrocytes. In a more recent study, Oh *et al*. [35] proposed a model involving glia-neuron interactions where activation of the astrocytic mechanosensitive channel TRPA1 could initiate extracellular glutamate release capable of altering the state of Glu-dependent ion channels in the neighboring neurons. In our study, we confirmed the dependance of the FUS-induced Ca^2+^ propagating waves over the extracellular glutamatergic signaling pathway (Fig. 6-IIIb). Follow-up studies should however be carried out to elucidate the respective roles of glial and neuronal cell populations in single-pulse FUS neurostimulation (Suppl. Discussion 3).

## 4 Conclusion

Overall, the exploration of a wide range of ultrasound frequencies appears to be a relevant tool both for the identification of the fundamental biophysical and neurobiological mechanisms underlying FUS neurostimulation and towards the development of therapeutic approaches offering acute spatial selectivity. The prospect of treatments in the megahertz-range is made possible via ultrasound delivery strategies based on intracranial transdural FUS implants, which could be particularly relevant in the context of chronic therapies. Results from our mechanistic study also suggest that megahertz-range FUS neurostimulation can promote the initiation of causal and targeted neural activities while limiting the risk of cavitation-induced damage, paving the way for a wide range of applications and fundamental research.

## Supporting information

Supplementary Video FUS 0.7

Supplementary Video FUS 2.2

Supplementary Video FUS 5.1

Supplementary Video FUS 8.1

Supplementary Video Mecha

Supplementary Video Flow off (control 4)

Supplementary Video Ca2+-free + Cd2+ + flow off

Supplementary Video Ca2+-free + Cd2+-free + flow off

Supplementary Video TTX + flow off

Supplementary Video Flow on (control 5)

Supplementary Video d-AP5 + flow

Supplementary Video Thapsigargin + flow

Supplementary Video Dantrolene + XeC + 2-APB + flow

Supplementary Video XeC + 2-APB + flow

Supplementary Video Dantrolene + flow

## 5 Acknowledgments

This project was supported by the French National Research Agency (ANR-16-TERC0017, ANR-21-CE19-0007 & ANR-21-CE19-0030), the American Focused Ultrasound Foundation (LabTAU, FUSF Center of Excellence). Additionally, this work was performed within the framework of the LABEX DEV WECAN (ANR-10-LABX-0061) and CORTEX (ANR-11-LABX-0042) of Université de Lyon, within the program “Investissements d’Avenir” (ANR-11-IDEX-0007) operated by the French National Research Agency (ANR). The work of the communities behind k-Wave, Blender and Python packages used throughout this work was integral to its completion. Finally, special gratitude goes to Nicola Kuczewski, Sandrine Parrot and Paul-Antoine Salin from the CRNL (*Centre de Recherche en Neuroscience de Lyon*) as well as Françoise Chavrier and Stefan Catheline from the LabTAU (Laboratory of Therapeutic Application of Ultrasound) who largely nourished the reflections around this study.

## 6 Methods

### 6.1 General investigation approach

All experiments were conducted on a model of *in vitro* human neural cells to form a 2D interconnected network. Cells were monitored using a Ca^2+^ fluorescence marker, and their responses to single-pulse FUS neurostimulation were studied on a fluorescence microscopy imaging platform equipped with a custom-made single-element focused ultrasound (FUS) transducer (Fig. 2-a). A wide range of ultrasound parameters were tested for their ability to elicit Ca^2+^ responses with a wide variety of spatiotemporal dynamics. A single-pulse FUS stimulation scheme was chosen on the basis of the groundwork established in TMS protocols for assessing a *motor threshold* (MT) [21], defined as the lowest stimulation level capable of eliciting a motor response with a single pulse. To keep the exploration of multiple FUS parameters manageable, the duration of the single FUS pulse was the only fixed parameter (400 µs). The feasibility of targeted single-pulse FUS stimulations was investigated by seeking minimal acoustic thresholds capable of inducing focal and causal Ca^2+^ mobilization over FUS frequencies ranging from 0.73 to 8.14 MHz. The exploration of parameters at such high frequencies was relevant in the context of the development of intracranial extradural FUS neurostimulation techniques. In addition to potential gains in the spatial selectivity of the stimuli provided when increasing frequencies, the use of various frequencies was expected to affect the predominance of frequency-dependent ultrasound-mediated biophysical processes, such as the probability of cavitation or the magnitude of FUS radiation force.

In addition to the proof of concept, stimulations in the presence of a fluorescent marker representative of mitochondrial cell activity were performed to assess cell tolerance to FUS stimulation (Fig. 2 and 3). Proper FUS targeting within the microscope field of view was ensured by applying stimulation pulses in cell-free Petri dishes filled with a solution of water and corn starch. This setup, combined with optical particle-tracking methods, enabled qualitative displacement maps to be established indicating the spatial distribution of mechanical stresses associated with each neurostimulation pulse. Analysis of the spatial distribution of these stresses and the emergence of Ca^2+^ responses in neural cultures were performed to shed light on the physical mechanisms underlying the observed FUS-tissue interactions. Hydrophone recordings of the acoustic field backscattered during stimulations were also acquired and analyzed using passive cavitation detection algorithms for comparison with spatial distribution analyses. In addition, numerical simulations have been carried out to estimate the acoustic and thermal perturbations induced in the Petri dish during FUS neurostimulation. These simulations are based on linearity assumptions that cannot fully describe all acoustic perturbations produced by FUS, for instance those produced indirectly by FUS through FUS-induced cavitation. These simulations were only indicative of acoustic and thermal disturbances induced by the direct acoustic contributions of the FUS field emitted by the transducer and its interactions with the acoustic domain (e.g., ultrasound attenuation by the cell subtract, ultrasound reflection by the petri dish). The spatiotemporal dynamics of intercellular Ca^2+^ responses propagating in the network after stimulations were studied with the aim of uncovering the signaling mechanisms mobilized by FUS neurostimulation.

A comparison was made with quasi-static singlecell mechanical stimulation using a patch-clamp micropipette (Fig. 3). The first interest was to consider a purely mechanical stimulation to determine whether FUS stimulation was mainly driven by mechanical or thermal effects. The second was to build on existing knowledge describing the neurobiology of neural responses obtained using this experimental framework [39, 52]. An extracellular flow (perfusion in the medium) was introduced (Fig. 4) to assess the involvement of extracellular messengers in the intercellular propagation of the elicited Ca^2+^ waves. Next, pharmacological blockade of Ca^2+^ and voltage-gated Na^+^ channels were tested to identify how ultrasound can evoke Ca^2+^ mobilization in the first place. Finally, pharmacological blockades were performed in conjunction with the extracellular flow to examine the role of specific signaling pathways (Fig. 5).

### 6.2 Single-pulse FUS generation

FUS stimulations were performed using a home-made single-element prototype consisting of a spherical ceramic transducer (C-21, Fuji Ceramics Corp.) with a diameter and focal length of 15 mm, embedded using resin (Stycast 2850FT, Loctite) in a SLS 3D-printed nylon casing (PA12, Xometry Europe). A function generator (Tektronix AFG1062, Tektronix Inc.) and a 50 dB power amplifier (Kalmus 150 RF, Amplifier Research Modular RF) were used to drive the transducer at its fundamental frequency (*f*_1_ = 0.73 MHz) or at its higher harmonics (*f*_3_ = 2.23 MHz, *f*_7_ = 5.19 MHz and *f*_11_ = 8.14 MHz). The values of these harmonics were obtained by measuring electrical input impedance of the transducer (Planar TR1300/1, Copper Mountain Technologies). The frequency and amplitude of the single-pulse FUS stimulation sequence (Fig. 2-a) (frequency, amplitude) were set with the function generator, using a continuous electrical sinusoidal signal. The duration of the pulse was defined by amplitude-modulation with a 400 µs square-wave pulse. Following previous works by our group [17], this duration was chosen on the basis of studies carried out in TMS protocols to define a *motor threshold* (MT) [21]. This modulation signal was itself triggered by the neuromonitoring platform.

### 6.3 Calcium fluorescence imaging

Calcium fluorescence recordings were performed using an inverted microscope (Eclipse Ti-S, Nikon Corp.) coupled to an ORCA-Fusion Hamamatsu Camera (Hamamatsu Photonics) operated at 10 frames per second. Fluorescence imaging was performed using a 494 nm excitation light (pE-300-W, CoolLED) coupled to a single-band emission filter transmitting at 516 nm (FITC-3540C-000, Semrock). The platform was placed in a faraday cage on an antivibration table (CleanBench Gimbal Piston, TMC) and equipped with micro-manipulators (MPC-200 - ROE-200 - MP-225, Sutter Instruments) for precise placement of patch-clamp pipettes and FUS transducers. For all stimulation conditions except FUS at 0.73 MHz, a 10 × optical magnification was applied (MRP00102, Nikon), providing a square 1.6 mm-wide field of view (FOV), with sufficient resolution to discern individual cells. Given the lateral dimension of the FUS focal spots, this optical magnification made it possible to assess both *focal* and *propagating* FUS-induced effects taking place in and around the target region for ultrasound frequencies as low as 2.23 MHz, as well as mechanical stimulation of single cells. On the other hand, for 0.73 MHz FUS stimulations, a 4 × lens (MRL00042, Nikon) providing a 3.9 mm FOV was preferred to match the larger focal spot size.

### 6.4 Combining FUS-sonication with calcium fluorescence microcopy imaging

The FUS transducer was attached to one of the micro-manipulator arms, enabling it to be positioned precisely in relation to the cell culture and microscope’s FOV. To ensure acoustic continuity between the transducer and the neural cell monolayer (+ bath solution), a 0.2% Agarose coupling gel (A9539 Sigma-Aldrich) was cast before the experiments in a 3D-printed PLA cone (333-60110, PLA Grey, Stratasys) (Fig. 1 and 2). Previous studies have estimated the sound velocity and acoustic impedance of the 0.2% agarose gel at 1503 m.s − − 1 & 1.52 MRayls, respectively [53, 54]. These estimates are close to those of the water-based cell media (1498 m.s − −1, 1.48 MRayls) allowing convenient transducer assembly while introducing limited amounts of acoustic reflections and aberrations. Single-pulse FUS stimulations were triggered by the fluorescence camera 10 s after the start of the fluorescence recording.

### 6.5 FUS targeting validation

Before stimulating the neural cells, proper FUS targeting was ensured by performing acoustic radiation force-driven displacement tests on an optical tissue phantom (Fig. 3). Based on the location of the displaced area, the location of the transducer assembly was adjusted using the micro-manipulator to be centered within the FOV. A PDMS-coated Petri dish was filled with 3 mL of a solution of water and cornstarch (Maïzena, Unilever). Once the starch had flowed and settled across the PDMS surface, the particles were illuminated laterally (OZM-A4516, Kern & Sohn) and used as an optical contrast agent to qualitatively assess the spatial distribution of FUS-induced mechanical stresses in the microscope FOV. As PDMS is optically transparent, surface displacements were highlighted by the addition of cornstarch acting as an optical contrast agent. The mobility of the contrasting particles also enabled us to highlight local disturbances such as those resulting from cavitation-induced micro-streaming. A Particle Image Velocimetry (PIV) algorithm, implemented in the openPIV Python library [55], was used to assess the relative displacement of these particles. The results of three to five image acquisitions performed on different regions of the Petri dish were combined via averaging to minimize the impact of contrast agent heterogeneities. With this method, FUS targeting could be assessed during the experiment on the basis of FUS-induced particle movements monitored on the live camera. The positioning of the transducer along *z* (ultrasound propagation axis) was adjusted by minimizing the diameter of the displaced region, ensuring that the top surface of the PDMS was correctly within the FUS focal area. Cumulative displacement maps were reconstructed by acquiring 100-frames movies of instantaneous particle displacements. The adjusted transducer position was recorded before using electronic micro-manipulators to move the transducer upwards. The Petri dish containing the optical phantom was then replaced by a Petri dish containing Human neural cells, and the transducer was moved down to its recorded position.

### 6.6 Passive cavitation detection of FUS-induced cavitation

In addition to the ultrasound transducer, a needle hydrophone (HNZ-1500, Onda corp.) was immersed in the cell medium to quantify the presence of FUS-induced cavitation events (Fig. 2 and 3). Introduced in biomedical applications of ultrasound to limit unsolicited damages [56] or ensure robust effects [57], methods for assessing cavitation levels following ultrasound sonications have been developed. Two classes of cavitation regimes have been identified [58]. While *stable* cavitation refers to a bubble state exhibiting weakly non-linear oscillations, bubble dynamics involving large oscillations and collapse are commonly referred to *inertial* cavitation. In the present study, inertial (IC) and stable (SC) cavitation indices were evaluated on the basis of the method described in [59]. While the IC gives an indication of the presence of broadband noise that may be associated to the emergence of bubble collapse events, the SC reflects the presence of quasi-linear oscillations at half the excitation frequency that are commonly recognized as characteristic of stable cavitation. IC is therefore expressed as the difference between the broadband noise level 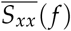 and a reference level *L*_ref_. These values were obtained by calculating the arithmetic mean of the hydrophone signal spectrum in the frequency range 0.1 − 9.0 MHz before and during ultrasound sonications. Obtained from a signal without ultrasound excitation, the reference level *L*_ref_ takes into account the background noise of the measurement. The occurrence of stable cavitation, on the other hand, was assessed from the difference between the average subharmonic noise and a reference emergent noise. These two values were obtained by averaging the spectrum amplitudes in windows centered around the subharmonic frequency defined as 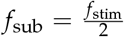. The width of these windows was arbitrarily chosen to be 6 kHz for subharmonic noise, to match the width scale of a potential peak while allowing some frequency variation. Conversely, emergent noise was estimated over a 600 kHz wide window not to be affected by a localized increase. wide window so as not to be affected by a local increase in noise.

### 6.7 Numerical modeling of FUS pressure field and FUS-induced heating

Numerical simulations (K-Wave, 1.4) [63] were performed to evaluate the acoustic field in the presence of the acoustic coupling cone (3D printed PLA plastic), the agarose gel, and the PDMS-coated Petri dish. The aCSF culture medium was acoustically simulated as water. Given geometry of the domain, the problem was reduced to a 2D axisymmetric configuration. Raster masks were created for each material domain to feed the medium properties with the values shown in Table 2. The spherical transducer was defined as a circular arc whose pressure magnitude was set, for each of the four FUS frequencies studied, on the basis of acoustic power measurements made separately using a radiation force balance (RFB-2000, Onda Corp.). Transducer surface pressures *p*_Tx_, were set as follows

**Table 2.** Acoustic and thermal material properties used for numerical simulations.

|  | Water [60] | Air [60] | PLA [61] | Agarose [53, 54] | PDMS [62] | Glass |
| --- | --- | --- | --- | --- | --- | --- |
| Sound speed $c$ [$\text{m.s}^{-1}$] | 1482.3 | 343 | 1250 | 1503 | 1054 | 5570 |
| Density $\rho$ [$\text{kg.m}^{-3}$] | 994.04 | 1.16 | 1240 | 1050 | 1038 | 2230 |
| Attenuation $\alpha_0$ [$\text{dB.cm}^{-1}.\text{MHz}^{-1}$] | 0.0022 | 1.907 | — | 0.014 | 0.315 | — |
| Attenuation non-linearity $b$ | 1 | 2 | — | 1 | 1.46 | — |
| Specific heat $c_t$ [$\text{J.kg}^{-1}.\text{K}^{-1}$] | 4178 | 1003.67 | — | $c_{t \text{ water}}$ | 1460 | 900 |
| Thermal conductivity $\kappa$ [$\text{W.m}^{-1}.\text{K}^{-1}$] | 0.6045 | 0.027 | — | $\kappa_{\text{water}}$ | 0.15 | 1.2 |

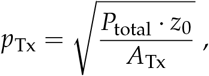

where*P*_total_ represents the total radiated acoustic power measured, *z*_0_ the characteristic acoustic impedance of the medium in which the transducer in immersed and *A*_Tx_ the transducer surface defined as a section of sphere such that

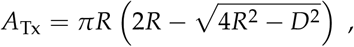

with *D* the transducer aperture and *R* its radius of curvature.

As the FUS stimulations consisted of a single, short and continuous pulse, *pulse average* acoustic quantities were reported. In order to convey information on the spatial distributions of these quantities, the *spatial average* and *spatial peak* were evaluated [64]. Given the diversity of ultrasound-derived mechanisms that may be involved, pressure, acoustic intensity and energy were reported. Thus *I*_sppa_ and *I*_sapa_ refer to the spatial peak pulse average and spatial average pulse average acoustic intensities in [W.cm^−2^]. *p*_sar_ and *p*_spr_ refer to the spatial average and peak RMS pressures, expressed in Pa. *E*_total_ refers to the total acoustic energy [J] radiated by the transducer, while *E*_focal spot_ refers to the acoustic energy deposited inside the focal spot. It should be noted that the focal spot areas used to evaluate these quantities were defined as the acoustic intensity full-width at half maximum (FWHM), i.e. at − 3 dB. Based on these acoustic simulations, FUS-induced heating was evaluated using the k-Wave implementation of the Bio Heat Transfer Equation (BHTE). Temperatures were computed at the PDMS-cell media interface, both in the attenuated domain of the PDMS itself and in the cell culture medium. As the culture medium had negligible acoustic attenuation (similar to that of water), negligible heating due to direct local absorption of FUS energy was expected. However, some heating was expected due to thermal diffusion of PDMS into the cell medium. With regard to the aforementioned acoustic quantities, *spatial average spatial peak* values were reported. Temporal descriptions were limited to *peak* values, associated with the time required to reach these maxima. Consequently, *spatial average temporal peak* temperature elevations were reported for PDMS and cell medium, *T*_PDMS satp_ and *T*_cells satp_ respectively, in conjunction with their *spatial peak* counterparts *T*_PDMS sptp_ and *T*_cells sptp_, all in [^°^C]. The times at which these temperature peaks occurred were expressed as *t*_T PDMS sptp_ and *t*_T cells sptp_ in [s]. Uncertainties relating to these quantities have been detailed in the statistical analysis section 6.15. Cell temperatures were evaluated on the grid points of the culture medium adjacent to the PDMS domain located at 109, 35, 15 & 10 µm from the water-PDMS interface respectively for simulations performed at 0.73 MHz, 2.23 MHz, 5.19 MHz and 8.14 MHz.

### 6.8 Quasi-static single-cell mechanical stimulations

Following the approach initiated by our team on large-scale neural models [65], quasi-static single-cell mechanical stimulations were performed with the aim of inducing highly targeted mechanical perturbations, in the absence of FUS-induced temperature elevations. A comparison was made with neural responses induced by FUS (Fig. 3). The first interest was to consider a purely mechanical stimulation to determine whether FUS stimulation was mainly driven by mechanical or thermal effects. The second was to build on existing knowledge describing the neurobiology of neural responses obtained using this experimental framework [39, 49, 66]. Borosilicate glass capillaries (1B150F-4, World Precision Instruments) were pulled and coupled to a patch clamp head-stage preamplifier (Axon Instruments CV-7B & MultiClamp 700B, Molecular Devices). An oscilloscope (PicoScope 3405D, Pico Technology) was used to monitor the electrical impedance of the pipette tip. This measurement, known as the *seal test*, provides information on pipette-cell contact and membrane integrity during the stimulation procedure. To achieve stimulation, the tip of the micropipette was first roughly positioned using a micro-manipulator in the center of the microscope’s FOV. Using a white light illumination and a 20× optical magnification objective, its location was then finely adjusted so that it rested on top of an identifiable cell body. In preparation for acquisition, the magnification was set back to 10 ×, the illumination switched to fluorescence excitation lighting and camera recording initiated. After acquisition of a 10 s baseline, the pipette was slowly lowered until the patch-clamp seal test signal indicated impedance values of the order of one giga Ohm, signaling correct contact between pipette and cell. Measurements were rejected if the seal test was lost (impedance measurements in the kilo Ohm range) before the end of the fluorescence acquisition, which could be attributed to cell membrane rupture. To match the chronology of FUS stimulation acquisitions, mechanical stimulation recordings were cut according to the timestamp of *giga seal* creation, in order to maintain a consistent 10 s pre-stimulation baseline followed by 30 s of post-stimulation recording.

### 6.9 In-vitro Human neural model preparation and comparison groups

In this study, all experiments were conducted on a human neural cell model in vitro. Immortalized human neural progenitors (ReNcell VM, Merck Millipore) were cultured and differentiated according to the protocol described in [67]. Cells were grown in 35 mm glass-bottom Petri dishes (Cellview, Greiner Bio-One) coated with a 2 mm thick layer of Polydimethylsiloxane, known as PDMS (Sylgard 184, Dow). At day 0, the surface of this PDMS substrate was first coated with laminin to promote cell adhesion, growth and differentiation. Then, after initial seeding at a density of 30 000 cells.cm^−2^, the progenitors were bathed for 3 days in artificial cerebrospinal fluid (aCSF) culture medium composed of ReNcell Maintenance Medium (SCM005, Merck Millipore), enriched with two *epidermal* and *basic epidermal* growth factors (GF0001 & GF003, Merck Millipore). At day 3, this enriched medium was removed and replaced by pure ReNcell maintenance medium for the remainder of the differentiation period. All neurostimulation experiments (FUS and quasi-static mechanical) were conducted at day 7 after the beginning of Human progenitor differentiation, the network being considered mature, appropriately confluent and structured with functional neurons and glial cells [1]. All the exploratory investigations to assess feasibility, repeatability, safety (cellular tolerance) and underlying biophysical and neurobiological mechanisms was carried out on 17 comparison groups, including: *1*. 3 groups to assess the safety of single-pulse FUS via the monitoring of mitochondrial activity(Fig. 2) and *2*. 14 groups where the mobilization of Ca^2+^ monitored to assess the characteristics of neural responses elicited by FUS stimulations vs those elicited by quasi-static mechanical ones (Fig. 3, 4 & 5).

The 3 groups evaluating the neural cell tolerance to single-pulse FUS were: *1*. two positive and negative control groups, **Viab. Sham** (control 1) and **Viab. Mecha. Death** (control 2) respectively with sham and destructive quasi-static mechanical stimulations and *2*. a **FUS 8.1 Viab**. group to assess neural cell viability after FUS. For all 3 groups, cell viability was assessed by adding 4 µL of a 200 µM mitochondria dye stock (Bio-Tracker 633 Red Mitochondria Dye, SCT137, Merck Millipore) to 2 mL of maintenance medium. Cell cultures were incubated with the viability dye for 30 min and then for the duration of the stimulation experiment. The composition of the aCSF extracellular media used throughout the study for the 13 groups in shown in Table 3. The 14 groups evaluating the FUS-evoked neural response and underlying mechanisms were as follows. Four groups, **FUS 0.7, FUS 2.2, FUS 5.2** and **FUS 8.1**, were tasked with studying the feasibility of single-pulse FUS neurostimulation at the 4 frequencies investigated (*f*_1_ = 0.73 MHz, *f*_3_ = 2.23 MHz, *f*_7_ = 5.19 MHz and *f*_11_ = 8.14 MHz). One group, **Mecha**. (control 3) was dedicated to evaluating the role of pure mechanical effects, as induced by quasi-static singlecell mechanical stimulation, in the responses evoked by FUS. Three groups, **Ca**^**2+**^**-free + Cd**^**2+**^ **+ flow off, Ca**^**2+**^**-free + flow off** and **TTX + flow off** were devoted to assessing the separate and combined roles of plasma Ca^2+^ channels and extracellular Ca^2+^ in gating intracellular processes potentially involved in FUS-induced responses at 8.14 MHz. They were directly compared to the **FUS 8.1** group which was extended to serve as a control group: **Flow off** (control 4). Six groups were finally formed to evaluate the influence of extracellular pathways on responses evoked by single-pulse FUS neurostimulation: *1*. a control group **Flow on** (control 5) to assess the effect of a moving extracellular medium on the spatial-temporal dynamics of neural responses evoked by FUS; *2*. a **d-AP5 + flow** group to specifically assess the role of the extracellular glutamate messenger, and *3*. **Thapsigargin + flow, Dantrolene + XeC + 2-APB + Flow, XeC + 2-APB + flow** and **Dantrolene + flow** groups to assess the separate and combined roles of intracellular Endoplasmic Reticulum (ER) ion pumps and channels in the chain of intracellular processes potentially involved in FUS-evoked responses, namely the SarcoEndoplasmic Reticulum Calcium ATPase (SERCA) pumps, Inositol 1.4.5-triphosphate Receptors (IP_3_R) and Ryanodine Receptors (RyR). For all of these 14 groups analyzing neural responses, a common cell preparation procedure was followed to enable assessment of calcium activity. At day 7, just prior to neurostimulation trials, cells were exposed for 30 min to a calcium-sensitive fluorescent marker (Fluo4 AM, AB241082, Abcam) in conjunction with a pluronic acid solution (Pluronic F-127, P3000MP, invitrogen) to permeabilize their membranes. After 30 min, the solution was discarded and replaced by 2 ml of pure ReNcell maintenance medium (Fig. 7).

**Table 3.** Composition of aCSF solutions used throughout the study. Control aCSF were used for all comparison groups at the exception of **Ca**^**2+**^**-free + Cd**^**2+**^ **+ flow off** and **Ca**^**2+**^**-free + flow off**.

| Concentrations [mM] | aCSF |  |  |
| --- | --- | --- | --- |
| | Control | $\text{Ca}^{2+}$-free + $\text{Cd}^{2+}$ | $\text{Ca}^{2+}$-free $\text{Cd}^{2+}$-free |
| NaCl | 125 | 125 | 125 |
| KCl | 2.5 | 2.5 | 2.5 |
| $\text{NaH}_2\text{PO}_4$ | 1.25 | 1.25 | 1.25 |
| $\text{NaHCO}_3$ | 25 | 25 | 25 |
| $\text{MgCl}_2$ | 1 | 3 | 3 |
| D+ glucose | 15 | 15 | 15 |
| Acide Ascorbique | 2 | 2 | 2 |
| Myo-Inositol | 3 | 3 | 3 |
| $\text{CaCl}_2$ | 2 | 0 | 0 |
| $\text{CdCl}_2$ | 0 | 0.1 | 0 |

### 6.10 Cellular tolerance to FUS neurostimulation

For the 3 groups evaluating cellular tolerance to single-pulse FUS neurostimulation, neural cell mitochondrial viability was followed by florescence microscopy imaging. Fluorescence images were acquired 2 min before FUS stimulation trials and every 2 min after stimulations over a 10 min period to assess the temporal dynamics of fluorescence staining. Positive and negative control groups were constituted with sham stimulations and destructive mechanical trials. The latter were performed with a patch clamp micropipette used to disrupt cell membrane integrity. Membrane disruption was monitored during the experiments using the seal test signal described in the quasi-static single-cell mechanical stimulation procedure. After initiating contact between the pipette and the cell validated by a giga seal, the pipette was slowly lowered using a micro-manipulator until the electrical impedance of the pipette returned to its pre-contact value. Mitochondrial viability after single-pulse FUS stimulations was compared with control groups (see **FUS 8.1 Viab**. vs **Viab. Sham** (control 1) and **Viab. Mecha. Death** (control 2)). Cell body regions of interests (ROIs) were manually defined for cells located in the region targeted by FUS stimulation. In the case of the destructive control group (see **Viab. Mecha. Death** (control 2)), a single ROI was defined on the cell subjected to the membrane disruption procedure described earlier in the methods. Fluorescence intensities were normalized to the baseline acquired at *t* = *t*_stim_ − 2 min.

### 6.11 Single-pulse FUS neurostimulation feasibility

The feasibility of single-pulse FUS neurostimulation was evaluated in terms of minimum FUS response threshold, causality, targeting selectivity and repeatability/success rate. Similar to the methodology used in clinical TMS protocols to assess a patient’s MT [21], a dose escalation strategy was followed by progressively increasing FUS pressure (and therefore FUS power, intensity and energy) to find the minimum ultrasound threshold to trigger an initial causal neural response with 400 µs single-pulse FUS stimulation. One of the first challenges was to determine under which FUS conditions a causal and reproducible neural response could be induced, while avoiding potential destructive effects (mechanical or thermal) induced by FUS. FUS stimulation were therefore repeated on distinct regions of a neural culture while progressively increasing pressure levels, up to 10 MPa spatial peak RMS or *p*_spr_ (Table 1). A minimum FUS threshold was sought for each of the four frequencies studied (0.73 MHz, 2.23 MHz, 5.19 MHz and 8.14 MHz). To reduce the number of variable ultrasound parameters and enable direct comparisons when exploring the effect of FUS frequencies on neurostimulation outcomes and underlying biophysical mechanisms, all investigations were conducted using the same FUS transducer driven at different harmonic frequencies (see **FUS 0.7, FUS 2.2, FUS 5.2** and **FUS 8.1** groups). The results of FUS neurostimulation were directly compared with those obtained with quasi-static single-cell mechanical stimulation (see **Mecha**. (control 3) group) (Fig 3 and 7). Within the range of ultrasound-evoked neural Ca^2+^ activity reported in the literature, particular attention was paid to a specific response regime in which spatially focal, temporally causal and subsequently propagating Ca^2+^ activity was initiated. We first identified four response regimes depending on the acoustic pressure level and frequency (Fig. 2): *1*. a first regime of *No measurable responses*, in which single-pulse stimuli at very-low pressures were unable to elicit measurable neural activity with the 10 fps acquisition configuration used in this study; *2*. A second regime of *Sparse responses*, in which increased pressures could result in elevated activity levels in individual cells throughout the microscope’s FOV. No clear signs of intercellular signaling were detectable at this stage. *3*. a third *Destructive* regime at higher acoustic pressures where the initiation of dispersed responses was possible with a drastic increase in the occurrence of large-scale cavitation events, often associated with cell damage and *4*. a fourth *Propagating* regime where discrete response initiation spatial distributions could be identified as a function of frequency. While low frequencies were associated to *Dispersed & Propagating* responses, increasing acoustic frequencies emerged as a viable means of reducing or eliminating the occurrence of destructive events while achieving acute spatial targeting and improved repeatability. This specific sub-regime was designated as *Focal & Propagating* Because of the increased reproducibility and control associated with the *Focal & Propagating* regime, the present study was designed to address the lack of descriptions of this specific response type. To assess causality, spatial selectivity, repeatability and describe the mechanisms underlying these FUS-evoked focal & propagating responses, two distinct periods were considered and classified as followed: *1. Immediate Focal* responses causally linked to the FUS stimulus and detected within 1 second of the onset of the single FUS pulse and *2. Delayed Propagating* responses appearing after 1 s and whose origin can be attributed to prior activation of neighboring cells during the *Immediate Focal* responses. Neural response causality was examined by monitoring cell activity before (at least 10 s before), during and after the single FUS pulse (at least 30 s after). Neurostimulation repeatability was assessed by quantifying a *neurostimulation success rate* (NSR) defined as the rate at which FUS could evoke an immediate focal neural response in at least one cluster of cells within the FUS focal spot area.

### 6.12 Analyses of the immediate focal Ca2+ responses

Once pressure thresholds capable of eliciting *propagating* responses had been found for each frequency, the influence of the single-pulse frequency on the spatial selectivity and distribution of Ca^2+^ responses was investigated. The accuracy of FUS targeting was assessed by analyzing the spatial distribution of *immediate* responses. In the *Dispersed* or *Focal & Propagating* class of FUS-elicited responses defined in Fig. 2-g, we identified that *immediate* Ca^2+^ activity was spatially structured in *clusters* immediately (≤1s) responding cells. Quantitative metrics in the name of *cluster diameters ϕ*, and *inter-cluster distances d* were established to characterize the spatial distribution of the neurostimulation *immediate* effects (≤ 1s) shown on Fig. 3-f-g. The metrics were also evaluated on displacement maps measured using an optical phantom. The method for the estimation of these maps is described in section 6.5.

Extraction of metrics were performed throughout manual annotation of *immediately responding cells* and *displacement* clusters with ellipses using the n-dimensional viewer Napari [68]. Note that in the *dispersed* regime associated to low frequencies (*f* ≤ 5 MHz), multiple clusters could simultaneously arise from a single FUS stimulation trial. When this happened, clusters were treated individually. All successful stimulation trials were pooled based on the stimulation frequency (**FUS 0.7, FUS 2.2, FUS 5.2 & FUS 8.1** groups) and 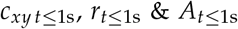, were defined respectively as the centroid, the radius and area of the immediately responding clusters. *Inter-cluster distances d* were computed based on clusters centers 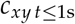 and *cluster diameters* were estimated to 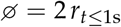.

The impact of the FUS frequency on cluster diameters and inter-cluster distances was then investigated. These metrics were also used to investigate the ultrasound biophysical mechanisms underlying FUS neurostimulation. It should be noted that for the sake of readability, these dimensions have been normalized by a coefficient 𝒩 defined as follows: *1. 𝒩*= *λ* for FUS neurostimulation, where *λ* refers to the ultrasound wavelength and *2. 𝒩* = 10 µm, corresponding to a characteristic neural cell body size for singe-cell quasi-static mechanical stimulation. This choice is explained by the fact that single-cell quasi-static mechanical stimulation enabled direct stimulation on the scale of a single body cell, while the effects induced by FUS were expected to be contained within the FUS focal spot. The characteristic lateral dimension of the − 3 dB focal spot of a spherical transducer can be approximated to

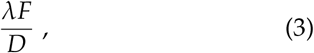

where *F* and *D* correspond to the focal length and aperture diameter respectively. As these dimensions were both equal to 15 mm in the presented setup, the FUS focal spot diameter thus was approximated to *λ*. Spatial distributions of simulated pressure levels, mechanical perturbations measured on the optical phantom and *focal* Ca^2+^ responses were compared across frequencies and confronted with evaluated passive cavitation indices to assess the preponderant physical mechanisms capable of inducing Ca^2+^ responses. Mechanical stresses *directly* induced by FUS pressure fields, such as radiation forces, were assessed by optical phantom measurements. Mechanical stresses indirectly induced by FUS pulses resulting from cavitation events were captured by optical phantom measurements and backscattered acoustic pressure recordings. Thermal effects directly induced by FUS were assessed by linear acousto-thermal numerical simulations. Although these methods alone cannot definitively dissociate the effects of mechanical and thermal perturbations on the triggering of the observed Ca^2+^ responses, a hypothesis without thermal elevation was explored by performing quasistatic single-cell mechanical stimulations.

### 6.13 Analyses of the delayed propagating Ca2+ responses

All analyses were conducted on *denoised fluorescence stacks* 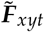 from which the baseline was subtracted (details on the processing of fluorescence images can be found in Supplementary Methods 1). As illustrated on Fig. 8 three distinct analyses were then performed on the filtered fluorescence image data. First, the temporal characteristics of the inter-cellular Ca^2+^ responses were assessed by automatic segmentation of neural cell bodies in the microscope’s 2D FOV, allowing extraction of fluorescence intensities for each individual cells (Fig. 8-b, 4 & 5). In combination with the aforementioned graphical analyses, cell segmentation maps were used to count the number *n*_cell_ of cell bodies in the FOV and quantify the percentages of cells *immediately* responding in the FUS focal spots 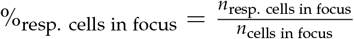, (Fig. 5-c). Peak cell fluorescence elevation rates, 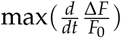, were calculated for neural cells located in the FOV as the maximum fluorescence variation over time. Second, the spatiotemporal characteristics of the intercellular Ca^2+^ response were assessed by constructing 3D graphs containing successive 2D spatial responses over time (Fig. 8-a). Single-cell response events represented in the 3D *xyt* space were fitted to models of intercellular signaling expansions. Based on these models, several metrics of the spatiotemporal dynamics of intercellular Ca^2+^ responses were established and evaluated. Expansion radii of the delayed propagating Ca^2+^ response, *r*(*t, φ*) were expressed using a diffusion-like model defined as

**Figure 8.**
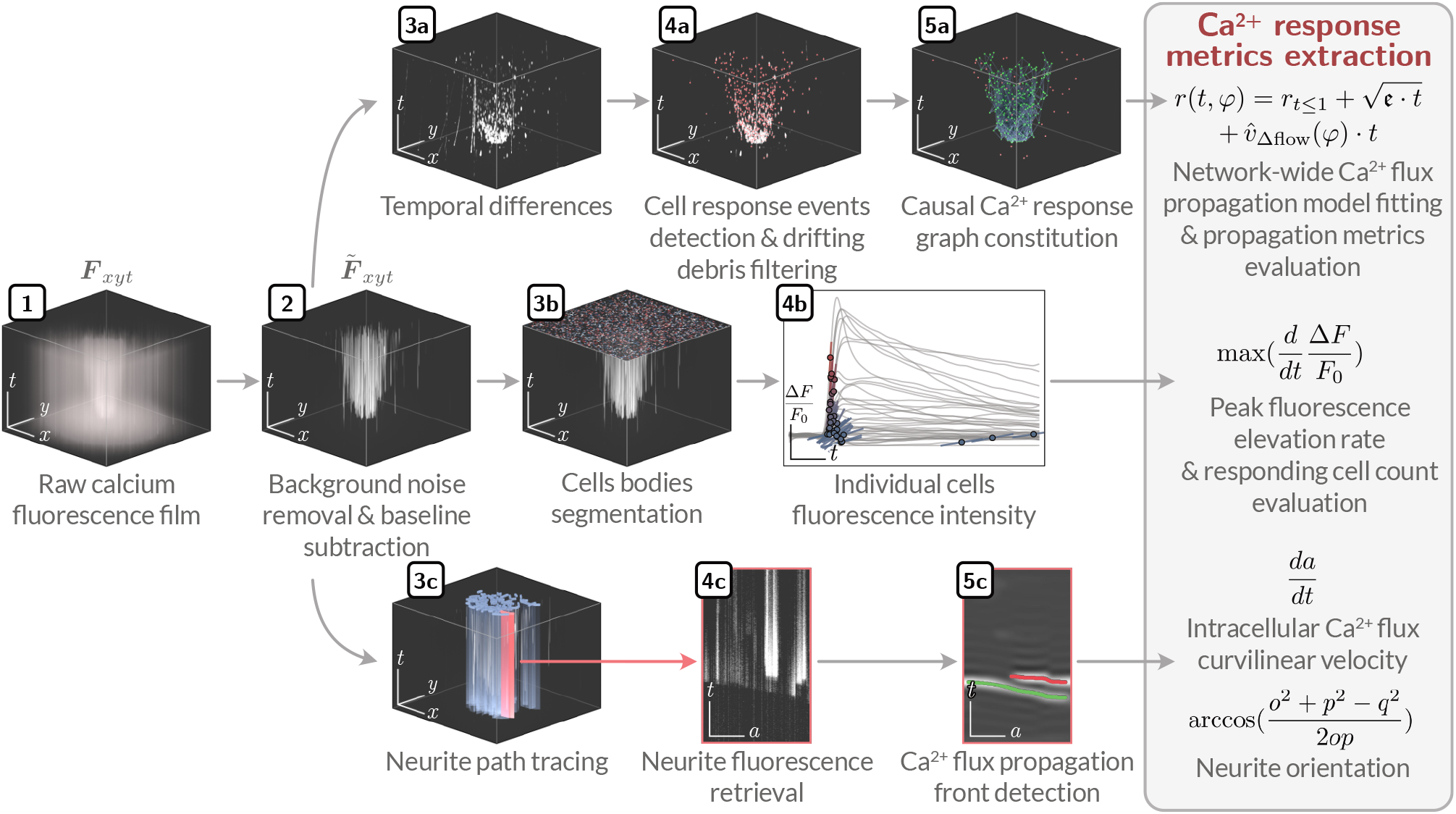
Overview of the Ca^2+^ fluorescence recording processing **pipeline** developed to extract metrics characterizing. **(section a)** the spatio-temporal dynamics of the network-wide response expansion, **(section b)** the temporal dynamics of individual cell fluorescence responses as well as **(section c)** the propagation velocity of Ca^2+^ waves along cultured neurites as a function of their orientation in the microscope FOV.

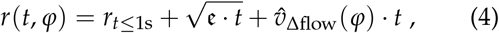

where 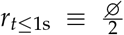 and 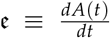 a constant with *φ* ∈ [0, 360]^°^ the angle of observation of radial propagation, *e* the surface expansion rate defined as a function of *A*(*t*), the cumulative response area covered by all neural cells that responded to neurostimulation, 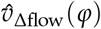 the radial drift velocity of the Ca^2+^ response in case of extracellular flow (Fig. 4 & 5). Without perfusion flow, the cell culture medium could be considered as static and expressions could be rewritten such that

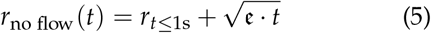

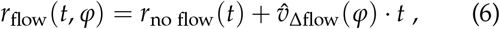

with *r*_no flow_(*t*) and *r*_flow_(*t, φ*) representing the expansion radii Ca^2+^ response with and without flow, respectively. Similar expressions could be derived for the expansion velocities, *v*_no flow_(*t*), *v*_flow_(*t, φ*), representing the velocities of the intercellular group of the diffusion-like response front (Fig. 4 and 5)

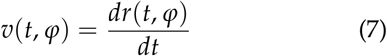

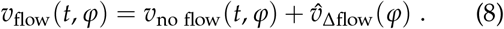

The Ca^2+^ response area expansion ratio mobilized in analyses involving pharmacological blockade (Fig. 5-b) was defined as the ratio of the response area at time *t* and the response area of the initial focal response

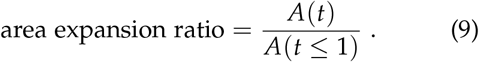

Finally, velocities of intracellular Ca^2+^ responses propagating along neurites were assessed by manual labeling of neurites using a series of segments allowing the automated detection of the Ca^2+^ propagation front (Fig. 8-c & Supplementary Methods 3). A 1D curvilinear referential frame was created for each neurite with a curvilinear abscissa *a*, defined by interpolating the coordinates in the *xy* cartesian reference frame of the neurite’s tortuous path. Next, the fluorescence history along *a* was constructed to perform the detection the propagating front of the intracellular Ca^2+^ response and estimate the velocities of the intracellular (intra-neurite) Ca^2+^ response. Each neurite annotation segment was located in the FOV and its azimuthal angle *φ* evaluated relative to the center 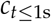 of the intercellular *immediate* response. As illustrated on Fig. 4-b, orientations *α* ∈ [0, 90]degree of neurites segments relative 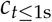 response were also defined. Empirical evaluation of the involvement of the intra- and extra-cellular pathways was achieved by performing the measurement of intracellular flux velocities on an *in silico* model for a purely extracellular cellular signaling pathway. The later was built by creating a synthetic fluorescence stack initially filled with zeros. Disk-shaped masks were then populated in each frame with radii based on equation 5 modeling network-wide calcium responses as a function of time. 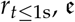 were derived from the median of the parameters fitted to FUS-induced responses from the **Flow off** (control 4) group. Neurite paths sampled from actual in vitro measurements were reemployed as to prevent biases introduced by variability in the sampled paths.

Detailed descriptions of the methods used to compute experimental and modeled parameters (e.g. filtering of fluorescence image, constructing numerical model of response velocities) can be found in the Data Analysis section of the Supplementary Methods.

### 6.14 Analyses of the neurobiological mechanisms

Two hypotheses of intra- and extra-cellular signaling pathways were formulated and tested to better understand the neurobiological mechanisms involved in the *propagative* neural responses evoked by single-pulse FUS neurostimulation. To assess the involvement of extracellular messengers such as glutamate - whose implication has been suggested in previous studies [35, 39] - FUS stimulations were reproduced in the presence of an extracellular perfusion. The hypothesis was that the spatial dynamic of any extra-cellular diffusion of chemical compounds involved in FUS-induced Ca^2+^ activities should be affected by an imposed extracellular flow in the cell culture medium, which is not expected for a purely intra-cellular signaling pathway. Extracellular flow was imposed by the addition of inlet and outlet tubes connected to a low-rate peristaltic pump (Peri-Star Pro 4LS, World Precision Instruments). These tubes were placed on either side of the transducer coupling cone, ensuring repeatable flow distribution across the microscope’s FOV. A temperature controller (TC-324C, Warner Instruments) was introduced before the inlet to limit excessive heat losses from the medium passing through the tube system. All investigations into neurobiological mechanisms were conducted using the FUS neurostimulation frequency configuration with the best performances in terms of control, repeatability and spatial selectivity (*f*_11_ = 8.14 MHz in this study). To evaluate the role of extracellular Ca^2+^ in FUS-induced *focal and propagating* Ca^2+^ responses, and to identify the entry pathways of mobilized Ca^2+^, FUS stimulations trials were conducted on cell cultures exposed to a blocker of plasma membrane Ca^2+^ channels. The study of the effect of this blockade on the FUS NSR provided a better understanding of the role of ion channels than the alternative hypothesis of sonoporation-induced ion exchange. Five minutes before stimulation trials, extracellular aCSF medium was replaced by aCSF in which CaCl_2_ was substituted by CdCl_2_. FUS stimulations were then performed in aCSFs with and without extracellular Ca^2+^ (Table 3). To assess the role of action potentials (APs) in the observed FUS-induced Ca^2+^ responses, the voltage-gated Na^+^ channel blocker TTX (Tetrodotoxin citrate, AB120055, Abcam) was introduced into Fluo4 rinsing medium at a concentration of 2 µM. The compound remained in contact with the cells during neurostimulation trials (see **Flow off** (control 4) vs **Ca**^**2+**^**-free + Cd**^**2+**^ **+ flow off, Ca**^**2+**^**-free + flow off** and **TTX + flow off** groups). Based on observations of the sensitivity of the propagating Ca^2+^ response to extracellular flow, we sought to identify the extracellular messengers involved in this FUS-evoked intercellular signaling process. Based on the works of Innocenti et al., Bazargani et al. or Oh et al. [35, 39, 69] and the results of quasi-static single-cell mechanical stimulations, glutamate signaling was identified as a first potential candidate. As NMDAR is the main glutamate receptor mediating Ca^2+^ entry into neurons and astrocytes, we looked for a change in response propagation in the presence of the NMDAR antagonist d-AP5. These trials conducted in the presence of extracellular flow enabled us to distinguish extracellular signaling from any intracellular pathway that might be involved. The NMDAR blocker (d-AP5, 18024662, Ficher Scientific) was therefore introduced into the extracellular medium at a concentration of 10 µM. A medium supplemented with d-AP5 was also used in the perfusion system for trials involving extracellular flows (see **Flow on** (control 5) vs **d-AP5 + flow** groups). To evaluate the involvement of Ca^2+^ amplification mechanisms, based on the discharge of intracellular Ca^2+^ stores, in FUS-elicited Ca^2+^activity, the effect of Ca^2+^ store depletion on the observed responses was studied. Depletion of intracellular Ca^2+^ stores was achieved via blockade of SarcoEndopblasmic Reticulum Calcium ATPase (SERCA) pumps using Thapsigargin (Thapsigargin, 586006, Merck Millipore) at a concentration of 10 µM. Finer identification of the signaling pathway involved in the discharge was investigated via blockade of the Ryanodine (RyR) and Inositol 1.4.5-triphosphate (IP3R) receptors. RyRs were inactivated using the RyR antagonist Dantrolene (Dantrolene sodium salt, D9175, Merck Millipore), while IP3R blockade was achieved using XeC (Xestospongin C, X2628, Merck Millipore) at 20 µM and 2-APB (2-APB, 100065, Merck Millipore) at 50 µM. In these blockade protocols, cells were exposed to the pharmacological treatments for a 20-min pre-incubation period prior to FUS stimulations trials. These two blockades were carried out both separately and in combination [70], and compared with the control group (see **Flow on** (control 5) vs **Thapsigargin + flow, Dantrolene + XeC + 2-APB + flow, XeC + 2-APB + flow** and **Dantrolene + flow** groups).

### 6.15 Statistical analyses and reporting

All statistical analyses were conducted using Prism software (GraphPad Prism 10, Insight Partners). Non-parametric Mann-Whitney tests were performed. Uncertainty ranges reported throughout this paper are expressed as median(*x*) 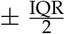, where IQR stands for the interquartile range defined as *Q*_3_ − *Q*_1_, the 75% and 25% quartiles respectively. In the case of acousto-thermal simulations results, quantities were evaluated based on the input voltages used for every stimulation trials. Reported values are expressed as median(*x*) 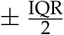 for each neural response.

## Supplementary Results *1* - On the correlation between FUS-induced thermal effects and the elicited Ca2+ responses

Although all above observations converged on predominant mechanical effects of FUS, the hypothesis of a FUS-induced thermal effect was also considered. Despite thermal effects induced by inertial cavitation could not be estimated, thermal effects directly induced by the absorption of the FUS field energy in the neural cell environment were numerically simulated. FUS-induced heating remained limited to the highly attenuating PDMS layer used as a mechanically compliant cell substrate (peaks Δ*T*_PDMS sptp_ = 0 − 18 ^°^C) (Table 1). Temperature elevations of neural cells were the result of thermal diffusion, according to the time delays observed between the heating peaks within the PDMS 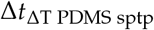 (equal to the 400 µs single-pulse FUS duration) and within cells 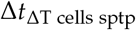 (tens of ms ≫ single-pulse FUS duration). No correlations were observed between the various FUS-induced heating (peaks Δ*T*_cells sptp_ = 0 − 7^°^C for *f* = 0.73 − 8.14 MHz) and the minimal thresholds of neurostimulation. Especially, when comparing configurations with a predominant direct effect of the FUS field (*f* ≥ 5.19 MHz), successful neurostimulations were ensured with half as much heating at 5.19 MHz (peak Δ*T*_cells sptp_ = 3 ^°^C) as at 8.14 MHz (peak Δ*T*_cells sptp_ = 7^°^C). This lack of correlation between temperature and neurostimulation was even more pronounced when comparing heating rates, ten times lower at 5.19 MHz (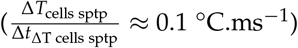) than at 8.14 MHz (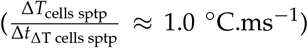). All these observations were not in favor of an intrinsic effect of FUS-induced heating in the observed single-pulse FUS neurostimulation phenomena.

### Supplementary Discussion *1* - Can single-pulse FUS neurostimulation be driven by thermal effects?

Unlike FUS neuromodulation, very few reports have been made on the possibility to evoke neurostimulation effects (according to Gavrilov’s definition [6]) via FUS-induced tissue heating. However, comparison with other techniques with similar spatial-temporal characteristics could help to estimate the potential impact of these effects. For instance, 1 ms long single-pulse infrared (IR) stimulations equating to energies of 2.8 mJ were reported to allow membrane depolarization capable of eliciting APs on wild type oocytes [71]. Although the cell model, the emitted energy nature, levels (2.8 mJ vs 0.5 J) and spatial distributions (400 µmwide laser pulse vs 180 µm-wide FUS focal spot) were not similar, reported energy pulse durations (1 ms single-pulse laser vs 400 µs single-pulse FUS) and absolute heating (Δ*T*≈ 10degreeC reached within oocytes vs Δ*T* = 3 − 7degreeC reached in neural cells) were comparable. In the laser stimulation study, however, the heating rate rather than the absolute heating has been highlighted as the main neurostimulation parameter. Heating rates were not comparable between the laser (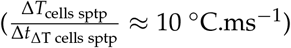) and the present FUS (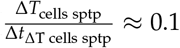 to 1.0 ^°^C.ms^−1^ from 5 to 8 MHz) studies. In addition, peak temperatures induced by laser were reached within 1 ms and by direct absorption of the optical energy in the cells. Peak temperatures induced by FUS were reached in the cells 7 to 23 ms after FUS, mostly by thermal diffusion effects from the highly absorbing PDMS-based cell subtract. Here, no clear correlation has been found between the FUS-induced absolute heating, the heating rate and the minimal conditions which could explain successful FUS neurostimulation. As a consequence, and considering above mentioned close similarities with quasi-static mechanical stimulations, thermal effects were not considered to be at the origin of the Ca^2+^ signaling induced by single-pulse FUS. On the other hand, the neuromodulatory effects of thermal heating have mostly been reported in terms of inhibitory rather than excitatory effects, with a reduction in the spiking activity (e.g. Δ*T* ≥ 30^°^C in astrocytes, [72, 73]). Any impact of FUS-induced heating on the neural response characteristics induced by single-pulse FUS should be further investigated to better understand any potential additional neuromodulatory effects occurring alongside the neurostimulatory effects that are the focus of the presented study.

### Supplementary Discussion *2* - Can the initiation of Ca2+ amplification not require an initial Ca2+ influx from the extracellular space?

A combination of the 3 known RyR isoforms (RyR1-3) is expressed in the mammalian brain, both in neurons and in non-neuronal cells (ref). CICR mechanisms induced by FUS could thus also involved RyR1 or RyR3 [74, 75]. An alternative hypothesis where the initiation of the Ca^2+^ amplification does not require extracellular Ca^2+^ entry into the cell can also be formulated with the RyR1 isoform considering descriptions made of depolarization-induced Ca^2+^ release (DICR) process (step **I bis**, a potential alternative to the hypothesis given in Fig. 6-**I**). In contrast to CICR, the DICR amplification was first described in skeletal muscle where communication between CaV1.1 and RyR1 takes place via a mechanical coupling, with conformational changes in CaV1.1 induced by cell membrane depolarization [76]. This mechanical interaction triggers the opening of RyR1 and subsequent release of intracellular Ca^2+^, without initial Ca^2+^ influx from the extracellular space [77, 78]. Although not described yet in neural cells, similar allosteric conformational transmission pathways could be involved in FUS neurostimulation. As an indirect effect of FUS on voltage-gated channels was not retained, a mechanically induced Ca^2+^ release (MICR) could be considered instead of a DICR mechanism. An initial direct mechanical action of FUS on a plasma ion channel (mechanically-sensitive, voltage-gated, ligand-gated) mechanically coupled to a RyR receptor could be an alternative mechanism for FUS-induced Ca^2+^ amplification.

### Supplementary Discussion 3 - Implication of astrocytes vs neurons in single-pulse FUS neurostimulation

The dynamics of Ca^2+^ flux observed show many similarities with descriptions of astrocytes responses to quasi-static mechanical stimulation [40]. In these studies, stimulations led to a slow (tens of µm.s^−1^) and omnidirectional Ca^2+^ flux propagating through the cellular networks. FUS-induced Ca^2+^ activities could then be mainly supported by astrocytes and gap junction-mediated intercellular signaling transmission. Given the importance of astrocytes for brain function [79], activation of astrocytic Ca^2+^ signaling pathway by single-pulse FUS neurostimulation could have important implications for future therapeutic FUS protocols. Other glial cells such as oligodendrocytes may also be involved in the Ca^2+^ signaling process observed, which could raise interest in developing other therapeutic mechanisms with FUS neurostimulation [80]. The main hypothesis of FUS-induced Ca^2+^ activity driven by non-neuronal cell types (astrocytes or oligodendrocytes) via gap-junction transmission was also supported by the fact that slow Ca^2+^ responses were unaffected by TTX blockade, indicating that APs and neurons were not the origin of the reported responses. Nevertheless, these observations could also remain compatible with neuronal cells expressing neuronal gap junctions [81, 82]. Indeed, neuronal gap junction-mediated Ca^2+^ waves have already been reported following mechanical stimulation of neuronal cells [66]. While they were affected by TTX, other studies have also reported spontaneous Ca^2+^ waves in differentiated cholinergic neuronal cells that were unaffected by TTX, similar to our observations [83]. In both cases, the involvement of glial-cell/glial-cell, glial-cell/neuron, and/or neuron/neuron gap junctions could allow messengers such Ca^2+^ or IP_3_ to mediate FUS-induced slow intercellular Ca^2+^ waves throughout these different cell types. Finally, confirmation of a slow, omnidirectional Ca^2+^ signaling process involving glial or neuronal cells expressing gap junctions does not preclude the existence of faster phenomena (*>*1 m.s^−1^) directly involving neuronal APs. Indeed, as pointed by Ross *et al*., widespread Ca^2+^ activities propagating through neurons can be driven by APs. In addition, our previous works conducted ex vivo on hippocampal brain slices (mouse model) using Micro-Electrode Array (MEA) neuromonitoring, showed that single-pulse FUS neurostimulation could elicit neuronal fiber volleys (FVs) and local field potentials (LFPs) [17]. However, if present in the current study, such fast Ca^2+^ signaling driven by APs was not detectable with our set-up optimized to study spatiotemporal dynamics at the cellular network scale. Follow-up studies could then be carried out while using alternative electrophysiological approaches (patch clamp, MEA) to elucidate the respective roles of glial and neuronal cell populations.

### Supplementary Methods 1 - Experimental data processing

For every stimulation trial, datasets were generated, included FUS transducer electrical driving commands before and after amplifications (function generator and amplifier outputs) and cavitation signals (hydrophone output) in csv format and the fluorescence imaging data (fluorescence camera output) recorded as tif films. This data were structured and processed using custom tools developed in Python. As the size of the fluorescence films reached several gigabytes, they were manipulated as dask arrays [84], limiting memory usage through lazy loading and speeding up operations by taking advantage of parallelized blocked algorithms. This structure allowed fluorescence films to be manipulated as 3D arrays containing both spatial and temporal dimensions, as illustrated in Suppl. Fig. 8. These objects will henceforth be referred to as fluorescence stacks or ***F***_*xyt*_. The general method of data analysis used in this study was to characterize of the focal and propagating Ca^2+^ responses elicited by FUS from fluorescence imaging data. All analyses were conducted on denoised fluorescence stacks from which the baseline was subtracted. Inconsistencies and fluctuations in the image background were first corrected by bandpass spatial filtering. The filter was designed on the basis of a modified Hann window such that

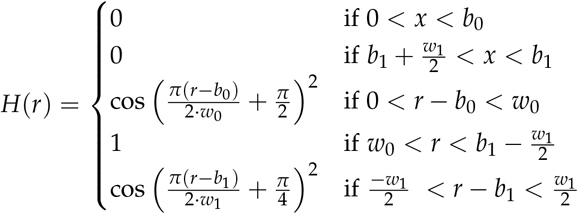

where *w*_0_ = 0.03, *w*_1_ = 0.35, *b*_1_ = 0.02 and *b*_1_ = 0.70. A 2D axisymmetric kernel was obtained by evaluating the function on a cylindrical coordinate meshgrid and the convolution of the fluorescence frames with the kernel was performed in the spatial frequency domain. Additionally, a fluorescence baseline selected as the frame at *t*_0_ − 5 s was subtracted from the whole stack as to eliminate optical perturbations emerging from stationary cell debris. Note that this background-free stack will be referred as 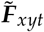.

### Supplementary Methods 1 - Assessment of the propagating response spatiotemporal dynamic

Intercellular *Focal & Propagating* Ca^2+^ responses elicited by 8.14 MHz FUS single-pulses were analyzed based on the constitution of a causal network graph. The latter was established after segmentation of individual cell Ca^2+^ response events as shown on Fig. 8-a. As to improve robustness to camera sensor noise and reduce the computational cost of subsequent operations, a spatially-binned copy of the stack was created through averaging using `dask’s coarsen` method with a 3 pixel-wide bin size along the *x* and *y* axes. Temporal fluorescence increase was then computed as the non-negative part of the difference of the left and right sides of a 1 s-long sliding hamming window. Using this representation, cell response events resulted in convex blob formations with spatial and temporal dimensions respectively analogous to the size of cell bodies and the duration taken to reach peak fluorescence intensity. Drifting fluorescent debris however created local perturbations that could be misclassified as cell responses. Hand tracing of these drifting paths was realized using Napari to generate masks applied to the stack to prevent false positives detection down the line. After filtering using a median filter, blob detection was performed using `pyclesperanto_prototype’s` GPU-accelerated 3D implementation of the Voronoi-Otsu-labelling algorithm with *outline* and *spot* sigmas of respectively 1 and 3. Using `pyclesperanto_prototype’s proximal_labels_to_igraph` method, a first physically-inaccurate cell response connectivity graph was established based on an arbitrary threshold distance of 60 pixels to limit the graph complexity. To ensure that causality was satisfied, graph edges were then screened based on the theoretical velocity needed to link two interconnected response event nodes *n*_*i*_, *n*_*j*_ defined as

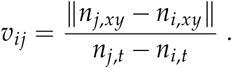

Given the slow nature of the Ca^2+^ waves we seek to study [49, 85], graph edges associated to theoretical velocities exceeding 100 µm.s^−1^ were discarded. Response nodes were then clustered into weakly connected component using Python’s igraph dedicated method. A collection of disconnected subgraphs was then obtained. Subgraphs representing stimulationevoked*propagating* Ca^2+^ responses were identified via the detection of *immediate response clusters*. As detailed in a previous section, individual cell response event nodes *n* occurring within a 1 second time-frame following mechanical or FUS stimulations were classified as being a potentially causal consequence of the stimulation. An arbitrary response subgraph *G* is classified as a propagating and stimulation-evoked intercellular Ca^2+^ response when *n*_immediate_ ∪ *G* and its number of constituting nodes exceeds 50. Based on this subgraph, a number of metrics characterizing network-wide calcium responses were established. Let us first define the center of the *focal immediate* response (*n*_immediate_, ∀ *n*_*k,t*_ ∈ ^≤^ 0 *t*_0_ ≤1) denoted as 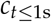. For the sake of readability in this section, variables will be written with subscripts indicating the node index *k* as well as its dimension in the *xyt* space. 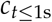 will therefore be referred as *c*_*xy*_. In practice, this location is evaluated as the centroid of the convex hull containing all subgraph nodes included in the [0, 1] s time interval. This spatial coordinate was used as a reference for the majority of metrics characterizing *focal & propagating* Ca^2+^ responses throughout this study.

While the distance of an individual cell response event to the intercellular response center is simply defined as ∥ *n*_*k,xy*_ − *c*_*xy*_∥, accurate estimation of their azimuthal angles required a preliminary correction step to account for the perfusion-induced drift. Using scikit-spatial’s best_fit function, the average drifting direction of the subgraph can be obtained as the vector *g*_*xyt*_ by fitting a line to the set of graph nodes. A perfusion-induced drift bias can then be expressed for each subgraph nodes such that *b*_*k,y*_ = *c*_*xy*_ + (*g*_*xyt*_ *n*_*k,t*_)_*xy*_. The azimuthal angle location *φ*^*k*^ of a node *k* was then evaluated as *φ*^*k*^ = arctan(*n*_*k,xy*_ − *b*_*k,xy*_). These parameters allowed the evaluation of Ca^2+^ responses expansion metrics defined in section 6.13.

### Supplementary Methods 2 Post-processing pipeline to detect intracellular propagating Ca2+ responses

#### Supplementary Methods 3 - Intra-cellular calcium flux velocity evaluation

Intra-cellular Ca^2+^ flux velocities were assessed by annotating neurite paths as collection of segments using the Napari viewer. Note that we used the term neurite to refer to all visibly elongated neural cell projections, as we did not performed a labelling test to assert their belonging to the *axon* or *dendrite* categories. As shown in Fig. 8-c, 2D matrices (time *t*, curvilinear abscissas of neurite *a*) of fluorescence intensities were retrieved for each hand-drawn paths. Ca^2+^ wavefronts propagating along the path were then detected following the signal processing pipeline detailed in Table 4. After proximity-based clustering detailed in the 8^th^ step of the pseudo algorithm, wavefronts were considered as valid if their length covered at least 95% of the neurite abscissas.

**Table 4.** Description of the post-processing pipeline developed for the detection of Ca^2+^ fluxes propagating along hand-segmented neurites paths as illustrated on Fig. 8-.

|  | Step Designation | Implementation |
| --- | --- | --- |
| 1 | Noise filtering filter | Weiner filter |
| 2 | Edge detection | Sobel filter<br>along the time axis |
| 3 | Noise filtering filter | Weiner filter |
| 4 | Local fluorescence<br>spike suppression | Morphological<br>(area opening) |
| 5 | Disregarding<br>fluorescence decrease | Negative value<br>clipping |
| 6 | Temporal<br>smoothing | Discarding high<br>frequency content |
| 7 | Local temporal<br>maxima detection | Peak detection using<br>`scipy.signal.find_peaks` |
| 8 | Proximity-based<br>clustering of<br>the detected peaks | Thresholds:<br>time: 1s,<br>abscissa: 3 pixels (2 $\mu\text{m}$) |

The velocity of their individual constituting segments was evaluated as 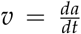. To elucidate the potential dependence of intracellular velocity on neurite orientation, the angle of hand-traced segment paths was evaluated. With *s*_0,*xy*_ and *s*_1,*xy*_ the start and end coordinates of a given segment path, we can define 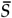 the center of this segment such that 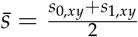. With *c*_*xy*_ the location of the intercellular response center. Orientations *α* of segments with respect to *c*_*xy*_ could be expressed as

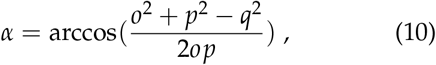

where 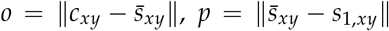 and *q* = ∥*s* ^1,*xy*^ − *c*^*xy*^∥.

As illustrated on Fig. 4-b, *α* angles were converted in degrees and wrapped within the [0 −90]°interval, where 0°and 90°correspond respectively to normal and tangential direction relative to the intercellular response center *c*_*xy*_.

### Supplementary Methods 4 - Analysis of responses with pharmacological blockade

The effects of the pharmacological treatments tested on the dynamics of response elicited by FUS at 8.14 MHz were evaluated at two levels. *1*. The temporal characteristics of Ca^2+^ mobilization at the individual cell level and *2*. the spatiotemporal attributes of network-wide Ca^2+^ responses. Based on the relation found in Fig. 3 between the theoretical lateral size of the FUS focal spot 3 and the size of the immediately elicited Ca^2+^ responses, we assessed the peak fluorescence elevation rate of individual cell body region of interests (ROIs) located at most half a wavelength from the intercellular response center.

Automatic segmentation of cell bodies was performed on the 50^th^ image of the fluorescence stack, corresponding to a 5 s baseline image before any stimulation. A pixel-binning operation was first realized (3 pixels bin size) before applying a contrast-limited adaptive histogram equalization (CLAHE) with a kernel size corresponding to the characteristic dimension of the cell bodies(`kernel_size=10, clip_limit=3e-2, nbins=2**12`). Actual cell body segmentation was then achieved by resizing the preprocessed image to the *x* and *y* pixel dimensions of the original stack and running `pyclesperanto_prototype’s voronoi_otsu_labeling` labelling algorithm (`spot_sigma=4.5, outline_sigma=3`). The centroids of the cell mask could then be evaluated and their distance from the center of the network-wide response computed. From the segmented masks, the number of responding cells was obtained and the fluorescence intensity signals for individual cells were extracted from background-free stack 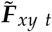 defined previously. These signals were then denoised using a second-order Butterworth low-pass filter (normalized critical frequency *W* = 0.2) and their time derivative was estimated using a forward difference scheme. Peak fluorescence elevation rates were obtained by taking the time derivative temporal peaks. Finally, outliers corresponding in most cases to fluorescent cell debris moving in and out of cell ROIs were filtered out, rejecting absolute peak elevation rates exceeding three time the standard deviation of the dataset.

The metrics established to characterize intercellular responses were based on the evaluation of their spatiotemporal envelope. The latter was constructed from the response graph defined in Fig. 8. The 3D *convex hull* of the graph’s vertices was computed using scipy’s dedicated ConvexHull function. For subsequent operations, this envelope was treated as a mesh object using Python’s trimesh package. Projections onto the *xy* plane of temporal slices 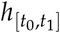 defined over the time interval [*t*_0_, *t*_1_] of the envelope mesh were used to evaluate the area, and centroids of the Ca^2+^ response during its propagation. The propagation of the response was assessed by means of an expansion ratio defined in equation 9.

The sensitivity of the response to extracellular flow was also evaluated by computing 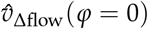 where *φ* = 0 corresponds to the direction of the flow. It should be noted that some FUS stimulation trial did not result in causal Ca^2+^ responses that would allow a response graph to be formed. The occurrence of these unsuccessful trials was reported as a Neurostimulation Success Rate (NSR).

## Footnotes

1 Ultrasound simulations performed for illustration purposes in CoperniFUS [8]. MRI data obtained from Schira *et al*. [9] with the authors’ permission. Graphics rendered using Blender.

## References

[1] Yuyu Song et al. “A Dynamic View of the Proteomic Landscape during Differentiation of ReNcell VM Cells, an Immortalized Human Neural Progenitor Line”. In: Scientific Data 6.1 (Mar. 2019), p. 190016. issn: 2052-4463. doi: 10.1038/sdata.2019.16.

[2] Valery L Feigin et al. “The Global Burden of Neurological Disorders: Translating Evidence into Policy”. In: The Lancet Neurology 19.3 (Mar. 2020), pp. 255– 265. issn: 14744422. doi: 10.1016/S1474-4422(19)30411-9.

[3] Joshua K. Wong et al. “Proceedings of the Ninth Annual Deep Brain Stimulation Think Tank: Advances in Cutting Edge Technologies, Artificial Intelligence, Neuromodulation, Neuroethics, Pain, Interventional Psychiatry, Epilepsy, and Traumatic Brain Injury”. In: Frontiers in Human Neuroscience 16 (Mar. 2022), p. 813387. issn: 1662-5161. doi: 10.3389/fnhum.2022.813387.

[4] Joshua K. Wong et al. “Proceedings of the 10th Annual Deep Brain Stimulation Think Tank: Advances in Cutting Edge Technologies, Artificial Intelligence, Neuromodulation, Neuroethics, Interventional Psychiatry, and Women in Neuromodulation”. In: Frontiers in Human Neuroscience 16 (Jan. 2023), p. 1084782. issn: 1662-5161. doi: 10.3389/fnhum.2022.1084782.

[5] Christine A. Edwards et al. “Neurostimulation Devices for the Treatment of Neurologic Disorders”. In: Mayo Clinic Proceedings 92.9 (Sept. 2017), pp. 1427– 1444. doi: 10.1016/j.mayocp.2017.05.005.

[6] Leonid Gavrilov. Use of Focused Ultrasound for Stimulation of Various Neural Structures. Nova Science Publishers. 2014. P.182. June 2014. isbn: 978-1-62948-9292.

[7] E. Newton Harvey. “The Effect of High Frequency Sound Waves on Heart Muscle and Other Irritable Tissues”. In: American Journal of Physiology-Legacy Content 91.1 (Dec. 1929), pp. 284–290. doi: 10.1152/ajplegacy.1929.91.1.284.

[8] Tom Aubier. Tomaubier/CoperniFUS: V0.1.2. Zenodo. June 2025. doi: 10.5281/ZENODO.15720551.

[9] Mark M. Schira et al. “HumanBrainAtlas: An in Vivo MRI Dataset for Detailed Segmentations”. In: Brain Structure and Function 228.8 (June 2023), pp. 1849– 1863. issn: 1863-2661. doi: 10.1007/s00429-023-02653-8.

[10] Lennart Verhagen et al. “Offline Impact of Transcranial Focused Ultrasound on Cortical Activation in Primates”. In: eLife 8 (Feb. 2019), e40541. issn: 2050-084X. doi: 10.7554/eLife.40541.

[11] Jean Francois Aubry. First Evidence of a Drastic and Sustained Reduction of Essential Tremor with Low Energy Transcranial Focused Ultrasound Neurostimulation: Clinical Report of 98% Tremor Reduction for More than 25 Minutes. Bethesda, MD, USA, Oct. 2022.

[12] R. Lalonde and J.W. Hunt. “Variable Frequency Field Conjugate Lenses for Ultrasound Hyperthermia”. In: IEEE Transactions on Ultrasonics, Ferroelectrics and Frequency Control 42.5 (Sept. 1995), pp. 825–831. issn: 0885-3010. doi: 10.1109/58.464838.

[13] Sergio Jiménez-Gambín et al. “Holograms to Focus Arbitrary Ultrasonic Fields through the Skull”. In: Physical Review Applied 12.1 (July 2019), p. 014016. issn: 2331-7019. doi: 10.1103/PhysRevApplied.12.014016.

[14] David Attali et al. “Deep Transcranial Ultrasound Stimulation Using Personalized Acoustic Metamaterials Improves Treatment-Resistant Depression in Humans”. In: Brain Stimulation (Apr. 2025), S1935861X25000993. issn: 1935861X. doi: 10.1016/j.brs.2025.04.018.

[15] Alexandre Carpentier et al. “Clinical Trial of BloodBrain Barrier Disruption by Pulsed Ultrasound”. In: Science Translational Medicine 8.343 (June 2016). doi: 10.1126/scitranslmed.aaf6086.

[16] Sara Cadoni et al. “Ectopic Expression of a Mechanosensitive Channel Confers Spatiotemporal Resolution to Ultrasound Stimulations of Neurons for Visual Restoration”. In: Nature Nanotechnology (Apr. 2023). issn: 1748-3387, 1748-3395. doi: 10.1038/s41565-023-01359-6.

[17] Ivan M. Suarez-Castellanos et al. “Spatio-Temporal Characterization of Causal Electrophysiological Activity Stimulated by Single Pulse Focused Ultrasound: An Ex Vivo Study on Hippocampal Brain Slices”. In: Journal of Neural Engineering 18.2 (Mar. 2021), p. 026022. doi: 10.1088/1741-2552/abdfb1.

[18] Eyal Weinreb and Elisha Moses. “Mechanistic Insights into Ultrasonic Neurostimulation of Disconnected Neurons Using Single Short Pulses”. In: Brain Stimulation (May 2022), S1935861X22000833. issn: 1935861X. doi: 10.1016/j.brs.2022.05.004.

[19] Byoung-Kyong Min et al. “Focused Ultrasound Modulates the Level of Cortical Neurotransmitters: Potential as a New Functional Brain Mapping Technique”. In: International Journal of Imaging Systems and Technology 21.2 (June 2011), pp. 232–240. issn: 08999457. doi: 10.1002/ima.20284.

[20] Shancheng Bao et al. “Personalized Depth-specific Neuromodulation of the Human Primary Motor Cortex via Ultrasound”. In: The Journal of Physiology (Feb. 2024), JP285613. issn: 0022-3751, 1469-7793. doi: 10.1113/JP285613.

[21] Carlo Miniussi, Walter Paulus, and Paolo M Rossini, eds. Transcranial Brain Stimulation. London, England: CRC Press, Oct. 2019.

[22] E. Newton Harvey and Alfred L. Loomis. “High Frequency Sound Waves of Small Intensity and Their Biological Effects”. In: Nature 121.3051 (Apr. 1928), pp. 622–624. issn: 0028-0836, 1476-4687. doi: 10.1038/121622a0.

[23] W. L. Nyborg and D. L. Miller. “Physical Mechanisms for Biological Effects of Ultrasound at Low-Intensity Levels”. In: Ultrasound Interactions in Biology and Medicine. Ed. by R. Millner, E. Rosenfeld, and U. Cobet. Boston, MA: Springer US, 1983, pp. 131– 138. isbn: 978-1-4684-8386-4 978-1-4684-8384-0. doi: 10.1007/978-1-4684-8384-0_18.

[24] William D. O’Brien. “Ultrasound–Biophysics Mechanisms”. In: Progress in Biophysics and Molecular Biology 93.1-3 (Jan. 2007), pp. 212–255. issn: 00796107. doi: 10.1016/j.pbiomolbio.2006.07.010.

[25] Michael Plaksin et al. “Thermal Transients Excite Neurons through Universal Intramembrane Mechanoelectrical Effects”. In: Physical Review X 8.1 (Mar. 2018), p. 011043. issn: 2160-3308. doi: 10.1103/PhysRevX.8.011043.

[26] Michael Plaksin, Shy Shoham, and Eitan Kimmel. “Intramembrane Cavitation as a Predictive Bio-Piezoelectric Mechanism for Ultrasonic Brain Stimulation”. In: Physical Review X 4.1 (Jan. 2014), p. 011004. issn: 2160-3308. doi: 10.1103/PhysRevX.4.011004.

[27] Martin Loynaz Prieto et al. “Dynamic Response of Model Lipid Membranes to Ultrasonic Radiation Force”. In: PLoS ONE 8.10 (Oct. 2013). Ed. by William Phillips, e77115. issn: 1932-6203. doi: 10.1371/journal.pone.0077115.

[28] Mike D. Menz et al. “Radiation Force as a Physical Mechanism for Ultrasonic Neurostimulation of the Ex Vivo Retina”. In: The Journal of Neuroscience 39.32 (Aug. 2019), pp. 6251–6264. issn: 0270-6474, 1529-2401. doi: 10.1523/JNEUROSCI.2394-18.2019.

[29] Jan Kubanek et al. “Ultrasound Modulates Ion Channel Currents”. In: Scientific Reports 6.1 (Apr. 2016), p. 24170. issn: 2045-2322. doi: 10.1038/srep24170.

[30] Sangjin Yoo et al. “Focused Ultrasound Excites Cortical Neurons via Mechanosensitive Calcium Accumulation and Ion Channel Amplification”. In: Nature Communications 13.1 (Jan. 2022), p. 493. issn: 2041-1723. doi: 10.1038/s41467-022-28040-1.

[31] Martin L Prieto and Merritt Maduke. “Toward an Ion-channel-centric Approach to Ultrasound Neuromodulation”. In: Current Opinion in Behavioral Sciences 56 (Apr. 2024), p. 101355. issn: 23521546. doi: 10.1016/j.cobeha.2024.101355.

[32] Pierre Pouget et al. “Neuronavigated Repetitive Transcranial Ultrasound Stimulation Induces LongLasting and Reversible Effects on Oculomotor Performance in Non-human Primates”. In: Frontiers in Physiology 11 (Aug. 2020). doi: 10.3389/fphys.2020.01042.

[33] Po Song Yang et al. “Transcranial Focused Ultrasound to the Thalamus Is Associated with Reduced Extracellular GABA Levels in Rats”. In: Neuropsychobiology 65.3 (2012), pp. 153–160. issn: 0302-282X, 1423-0224. doi: 10.1159/000336001.

[34] Siti N. Yaakub et al. “Transcranial Focused Ultrasound-Mediated Neurochemical and Functional Connectivity Changes in Deep Cortical Regions in Humans”. In: Nature Communications 14.1 (Sept. 2023), p. 5318. issn: 2041-1723. doi: 10.1038/s41467-023-40998-0.

[35] Soo-Jin Oh et al. “Ultrasonic Neuromodulation via Astrocytic TRPA1”. In: Current Biology 29.20 (Oct. 2019), 3386–3401.e8. doi: 10.1016/j.cub.2019.08.021.

[36] Malachy Newman et al. “Ultrasound Modulates Calcium Activity in Cultured Neurons, Glial Cells, Endothelial Cells and Pericytes”. In: Ultrasound in Medicine & Biology (Dec. 2023), S0301562923003642. issn: 03015629. doi: 10.1016/j.ultrasmedbio.2023.11.004.

[37] Benjamin Clennell et al. “Ultrasound Modulates Neuronal Potassium Currents via Ionotropic Glutamate Receptors”. In: Brain Stimulation 16.2 (Mar. 2023), pp. 540–552. issn: 1935861X. doi: 10.1016/j.brs.2023.01.1674.

[38] EncyclopediaofMathBernoulli. Bernoulli Random Walk -Encyclopedia of Mathematics. [50] https://encyclopediaofmath.org/wiki/Bernoulli_random_walk. 2025.

[39] Barbara Innocenti, Vladimir Parpura, and Philip G. Haydon. “Imaging Extracellular Waves of Glutamate during Calcium Signaling in Cultured Astrocytes”. In: The Journal of Neuroscience 20.5 (Mar. 2000), pp. 1800–1808. doi: 10.1523/jneurosci.20-05-01800.2000.

[40] Andrew Charles. “Intercellular Calcium Waves in Glia”. In: Glia 24.1 (Sept. 1998), pp. 39–49. issn: 0894-1491, 1098-1136. doi: 10.1002/(SICI)1098-1136(199809)24:1<39::AID-GLIA5>3.0.CO;2-W.

[41] William N. Ross. “Understanding Calcium Waves and Sparks in Central Neurons”. In: Nature Reviews Neuroscience 13.3 (Mar. 2012), pp. 157–168. issn: 1471003X, 1471-0048. doi: 10.1038/nrn3168.

[42] Nicholas E. Karagas and Kartik Venkatachalam. “Roles for the Endoplasmic Reticulum in Regulation of Neuronal Calcium Homeostasis”. In: Cells 8.10 (Oct. 2019), p. 1232. issn: 2073-4409. doi: 10.3390/cells8101232.

[43] Hermes A. S. Kamimura et al. “Ultrasound Neuromodulation: Mechanisms and the Potential of Multimodal Stimulation for Neuronal Function Assess-ment”. In: Frontiers in Physics 8 (May 2020), p. 150. issn: 2296-424X. doi: 10.3389/fphy.2020.00150.

[44] Fenfang Li et al. “Dynamics and Mechanisms of Intracellular Calcium Waves Elicited by Tandem Bubble-Induced Jetting Flow”. In: Proceedings of the National Academy of Sciences 115.3 (Jan. 2018). issn: 0027-8424, 1091-6490. doi: 10.1073/pnas.1713905115.

[45] Elisabetta Sassaroli and Natalia Vykhodtseva. “Acoustic Neuromodulation from a Basic Science Prospective”. In: Journal of Therapeutic Ultrasound 4.1 (May 2016). doi: 10.1186/s40349-016-0061-z.

[46] AlexanderG. Petrov et al. “Flexoelectric Effects in Model and Native Membranes Containing Ion Channels”. In: European Biophysics Journal 22.4 (Oct. 1993). issn: 0175-7571, 1432-1017. doi: 10.1007/BF00180263.

[47] William J. Tyler. “The Mechanobiology of Brain Function”. In: Nature Reviews Neuroscience 13.12 (Dec. 2012), pp. 867–878. issn: 1471-003X, 1471-0048. doi: 10.1038/nrn3383.

[48] Benjamin M. Gaub et al. “Neurons Differentiate Magnitude and Location of Mechanical Stimuli”. In: Proceedings of the National Academy of Sciences 117.2 (Jan. 2020), pp. 848–856. issn: 0027-8424, 1091-6490. doi: 10.1073/pnas.1909933117.

[49] Luc Leybaert and Michael J. Sanderson. “Intercellular Ca^2+^Waves: Mechanisms and Function”. In: Physiological Reviews 92.3 (July 2012), pp. 1359–1392. issn: 0031-9333, 1522-1210. doi: 10.1152/physrev.00029.2011.

[50] Satoko Soga-Sakakibara et al. “Calcium Dependence of the Priming, Activation and Inactivation of Ryanodine Receptors in Frog Motor Nerve Terminals”. In: European Journal of Neuroscience 32.6 (Sept. 2010), pp. 948–962. issn: 0953-816X, 1460-9568. doi: 10.1111/j.1460-9568.2010.07381.x.

[51] Nancy L. Allbritton, Tobias Meyer, and Lubert Stryer. “Range of Messenger Action of Calcium Ion and Inositol 1,4,5-Trisphosphate”. In: Science 258.5089 (Dec. 1992), pp. 1812–1815. issn: 0036-8075, 1095-9203. doi: 10.1126/science.1465619.

[52] Andrew C. Charles et al. “Intercellular Signaling in Glial Cells: Calcium Waves and Oscillations in Response to Mechanical Stimulation and Glutamate”. In: Neuron 6.6 (June 1991), pp. 983–992. doi: 10.1016/0896-6273(91)90238-u.

[53] S. Yang et al. “High-Intensity Focused Ultrasound Ablation: An in Vitro Agarose Gel Model”. In: International Journal of Clinical and Experimental Medicine 10 (Nov. 2017), pp. 15302–15308.

[54] Martin O. Culjat et al. “A Review of Tissue Substitutes for Ultrasound Imaging”. In: Ultrasound in Medicine & Biology 36.6 (June 2010), pp. 861–873. doi: 10.1016/j.ultrasmedbio.2010.02.012.

[55] Alex Liberzon et al. OpenPIV/Openpiv-Python: OpenPIV - Python (v0.22.2) with a New Extended Search PIV Grid Option. 2020. doi: 10.5281/ZENODO.3930343.

[56] A. Novell et al. “A New Safety Index Based on Intrapulse Monitoring of Ultra-Harmonic Cavitation during Ultrasound-Induced Blood-Brain Barrier Opening Procedures”. In: Scientific Reports 10.1 (June 2020), p. 10088. issn: 2045-2322. doi: 10.1038/s41598-020-66994-8.

[57] Abbas Sabraoui et al. “Feedback Loop Process to Control Acoustic Cavitation”. In: Ultrasonics Sonochemistry 18.2 (Mar. 2011), pp. 589–594. issn: 13504177. doi: 10.1016/j.ultsonch.2010.07.011.

[58] RE Apfel. “ACOUSTIC CAVITATION: A POSSIBLE CONSEQUENCE OF BIOMEDICAL USES OF ULTRASOUND”. In: (1982).

[59] Corentin Cornu et al. “Ultrafast Monitoring and Control of Subharmonic Emissions of an Unseeded Bubble Cloud during Pulsed Sonication”. In: Ultrasonics Sonochemistry 42 (Apr. 2018), pp. 697–703. doi: 10.1016/j.ultsonch.2017.12.026.

[60] IT’IS Foundation. Tissue Properties Database V4.1. 2022. doi: 10.13099/VIP21000-04-1.

[61] Depeng Ma et al. “Mechanical and Medical Imaging Properties of 3D-printed Materials as Tissue Equivalent Materials”. In: Journal of Applied Clinical Medical Physics 23.2 (Feb. 2022), e13495. issn: 1526-9914, 1526-9914. doi: 10.1002/acm2.13495.

[62] A. Cafarelli et al. “Tuning Acoustic and Mechanical Properties of Materials for Ultrasound Phantoms and Smart Substrates for Cell Cultures”. In: Acta Biomaterialia 49 (Feb. 2017), pp. 368–378. issn: 17427061. doi: 10.1016/j.actbio.2016.11.049.

[63] Bradley E. Treeby and B. T. Cox. “K-Wave: MATLAB Toolbox for the Simulation and Reconstruction of Photoacoustic Wave Fields”. In: Journal of Biomedical Optics 15.2 (2010), p. 021314. issn: 10833668. doi: 10.1117/1.3360308.

[64] Roy C. Preston et al. Output Measurements for Medical Ultrasound. Ed. by Roy C. Preston. London: Springer London, 1991. isbn: 978-1-4471-1885-5 978-1-4471-1883-1. doi: 10.1007/978-1-4471-1883-1.

[65] Jérémy Vion-Bailly et al. “Neurostimulation Success Rate of Repetitive-pulse Focused Ultrasound in an in Vivo Giant Axon Model: An Acoustic Parametric Study”. In: Medical Physics 49.1 (Jan. 2022), pp. 682– 701. issn: 0094-2405, 2473-4209. doi: 10.1002/mp.15358.

[66] Andrew C. Charles, Susheel K. Kodali, and Rachel F. Tyndale. “Intercellular Calcium Waves in Neurons”. In: Molecular and Cellular Neuroscience 7.5 (May 1996), pp. 337–353. issn: 10447431. doi: 10.1006/mcne.1996.0025.

[67] Roberta Donato et al. “Differential Development of Neuronal Physiological Responsiveness in Two Human Neural Stem Cell Lines”. In: BMC Neuro-science 8.1 (Dec. 2007), p. 36. issn: 1471-2202. doi: 10.1186/1471-2202-8-36.

[68] Jannis Ahlers et al. Napari: A Multi-Dimensional Image Viewer for Python. Zenodo. July 2023. doi: 10.5281/ZENODO.3555620.

[69] Narges Bazargani and David Attwell. “Astrocyte Calcium Signaling: The Third Wave”. In: Nature Neu-roscience 19.2 (Feb. 2016), pp. 182–189. issn: 1097-6256, 1546-1726. doi: 10.1038/nn.4201.

[70] Jessica Gambardella et al. “The Discovery and Development of IP3 Receptor Modulators: An Update”. In: Expert Opinion on Drug Discovery 16.6 (June 2021), pp. 709–718. issn: 1746-0441, 1746-045X. doi: 10.1080/17460441.2021.1858792.

[71] Mikhail G. Shapiro et al. “Infrared Light Excites Cells by Changing Their Electrical Capacitance”. In: Nature Communications 3.1 (Mar. 2012), p. 736. issn: 2041-1723. doi: 10.1038/ncomms1742.

[72] Carola G. Schipke et al. “Temperature and Nitric Oxide Control Spontaneous Calcium Transients in Astrocytes”. In: Cell Calcium 43.3 (Mar. 2008), pp. 285– 295. issn: 01434160. doi: 10.1016/j.ceca.2007.06.002.

[73] Niko Komin et al. “Multiscale Modeling Indicates That Temperature Dependent [Ca ^2+^] ^i^Spiking in Astrocytes Is Quantitatively Consistent with Modulated SERCA Activity”. In: Neural Plasticity 2015 (2015), pp. 1–15. issn: 2090-5904, 1687-5443. doi: 10.1155/2015/683490.

[74] Fumiaki Mori et al. “Developmental Changes in Expression of the Three Ryanodine Receptor mR-NAs in the Mouse Brain”. In: Neuroscience Letters 285.1 (May 2000), pp. 57–60. issn: 03043940. doi: 10.1016/S0304-3940(00)01046-6.

[75] Rodrigo Torres and Cecilia Hidalgo. “Subcellular Localization and Transcriptional Regulation of Brain Ryanodine Receptors. Functional Implications”. In: Cell Calcium 116 (Dec. 2023), p. 102821. issn: 01434160. doi: 10.1016/j.ceca.2023.102821.

[76] Hiroshi Takeshima et al. “Primary Structure and Expression from Complementary DNA of Skeletal Muscle Ryanodine Receptor”. In: Nature 339.6224 (June 1989), pp. 439–445. issn: 0028-0836, 1476-4687. doi: 10.1038/339439a0.

[77] Kellie A. Woll and Filip Van Petegem. “Calcium-Release Channels: Structure and Function of IP<sub>3</sub> Receptors and Ryanodine Receptors”. In: Physiological Reviews 102.1 (Jan. 2022), pp. 209–268. issn: 0031-9333, 1522-1210. doi: 10.1152/physrev.00033.2020.

[78] BA Block et al. “Structural Evidence for Direct Interaction between the Molecular Components of the Transverse Tubule/Sarcoplasmic Reticulum Junction in Skeletal Muscle.” In: The Journal of cell biology 107.6 (Dec. 1988), pp. 2587–2600. issn: 0021-9525, 1540-8140. doi: 10.1083/jcb.107.6.2587.

[79] Jérôme Ribot et al. “Astrocytes Close the Mouse Critical Period for Visual Plasticity”. In: Science 373.6550 (July 2021), pp. 77–81. issn: 0036-8075, 1095-9203. doi: 10.1126/science.abf5273.

[80] Manasi Iyer et al. “Oligodendrocyte Calcium Signaling Promotes Actin-Dependent Myelin Sheath Extension”. In: Nature Communications 15.1 (Jan. 2024), p. 265. issn: 2041-1723. doi: 10.1038/s41467-023-44238-3.

[81] Goran Söhl, Stephan Maxeiner, and Klaus Willecke. “Expression and Functions of Neuronal Gap Junctions”. In: Nature Reviews Neuroscience 6.3 (Mar. 2005), pp. 191–200. issn: 1471-003X, 1471-0048. doi: 10.1038/nrn1627.

[82] Rafael Gutiérrez. “Gap Junctions in the Brain: Hardwired but Functionally Versatile”. In: The Neuroscientist 29.5 (Oct. 2023), pp. 554–568. issn: 1073-8584, 1089-4098. doi: 10.1177/10738584221120804.

[83] Nishani T. Hettiarachchi et al. “Gap Junction-Mediated Spontaneous Ca2+ Waves in Differentiated Cholinergic SN56 Cells”. In: Biochemical and Biophysical Research Communications 397.3 (July 2010), pp. 564–568. issn: 0006291X. doi: 10.1016/j.bbrc.2010.05.159.

[84] Matthew Rocklin. “Dask: Parallel Computation with Blocked Algorithms and Task Scheduling”. In: Python in Science Conference. Austin, Texas, 2015, pp. 126–132. doi: 10.25080/Majora-7b98e3ed-013.

[85] Lionel F. Jaffe. “Fast Calcium Waves”. In: Cell Calcium 48.2-3 (Aug. 2010), pp. 102–113. issn: 01434160. doi: 10.1016/j.ceca.2010.08.007.

